# Condensin II collaborates with cohesin to establish and maintain interphase chromosome territories

**DOI:** 10.1101/2025.07.25.666893

**Authors:** Takao Ono, Masatoshi Takagi, Hideyuki Tanabe, Tomoko Fujita, Noriko Saitoh, Akatsuki Kimura, Tatsuya Hirano

**Affiliations:** Chromosome Dynamics Laboratory, RIKEN, Saitama, Japan; Cellular Dynamics Laboratory, RIKEN, Saitama, Japan; Research Center for Integrative Evolutionary Science, The Graduate University for Advanced Studies, SOKENDAI, Kanagawa, Japan; Division of Cancer Biology, The Cancer Institute of JFCR, Tokyo, Japan; Cell Architecture Laboratory, Department of Chromosome Science, National Institute of Genetics, Shizuoka, Japan; Present address: Laboratory for Cell Function Dynamics, RIKEN Center for Brain Science, Saitama, Japan

## Abstract

Despite the well-established role of condensin II in mitotic chromosome assembly, its function in interphase chromosome organization remains poorly understood. Here, we applied multiscale fluorescence in situ hybridization (FISH) techniques to human cell lines engineered for single or double depletion of condensin II and cohesin, and examined their functional interplay at two distinct stages of the cell cycle. Our results demonstrate that a functional “handover” from condensin II to cohesin during the mitosis-to-G1 transition is critical for establishing chromosome territories (CTs) in the newly assembling nucleus. During G2 phase, condensin II and cohesin cooperate to maintain global CT morphology, although they act at different genomic scales. Strikingly, double depletion of both complexes causes CTs to collapse and accumulate abnormally at the nucleolar periphery. Based on these findings, we will discuss how the condensin and cohesin complexes act in an orderly and cooperative manner to orchestrate chromatin dynamics across genomic scales, thereby supporting higher-order chromosome organization throughout the cell cycle.

## Introduction

Chromosomes undergo a series of dynamic conformational changes throughout the cell cycle. One of the most striking events occurs at the onset of mitosis, when the amorphous mass of interphase chromatin fibers is transformed into a discrete set of rod-shaped mitotic chromosomes (Batty and Gerlich, 2019; Paulson et al., 2021; Hirano, 2025). This process, known as mitotic chromosome assembly, is an essential prerequisite for the faithful segregation of sister chromatids in subsequent anaphase, and is mediated by two large protein complexes, condensin I and condensin II. The two condensin complexes share a common pair of structural maintenance of chromosomes (SMC) core subunits, SMC2 and SMC4, but differ in their non-SMC regulatory subunits: condensin I contains CAP-D2,-G, and-H, whereas condensin II contains CAP-D3,-G2, and-H2 (Ono et al., 2003). Notably, the two complexes exhibit distinct spatiotemporal dynamics during the cell cycle (Hirota et al., 2004; Ono et al., 2004; Gerlich et al., 2006). In brief, condensin II localizes to the nucleus during interphase and participates in the early stages of mitotic chromosome assembly in prophase. In contrast, condensin I is sequestered in the cytoplasm from interphase through prophase and gains access to chromosomes only after nuclear envelope breakdown (NEBD) in prometaphase. In metaphase, both condensin I and condensin II are enriched along the longitudinal axes of sister chromatids (Ono et al., 2003; Green et al., 2012), where they organize chromatin loops of different sizes (Gibcus et al., 2018; Samejima et al., 2025).

The next major transition occurs from mitotic telophase to early G1, when rod-shaped chromosomes are transformed back into spherical chromosome territories (CTs) in the newly reassembling nucleus (Fig. 1 A)(Cremer and Cremer, 2010). During this M-to-G1 transition, condensin I first dissociates from chromosomes and is exported out of the nucleus, followed by the dissociation of condensin II, which remains within the nucleus. Recent studies have reported that condensin II depletion during this transition impairs post-mitotic centromere dispersion in both animal and plant cells (Hoencamp et al., 2021; Sakamoto et al., 2022). Moreover, high-throughput chromosome conformation capture (Hi-C) analysis revealed that, under such conditions, chromosomes retain a parallel arrangement of their arms, reminiscent of the so-called Rabl configuration (Hoencamp et al., 2021). These findings indicate that the loss or impairment of condensin II during mitosis results in defects in post-mitotic chromosome organization. Notably, cohesin replaces condensin II during this transition (Fig. 1 B)(Abramo et al., 2019; Brunner et al., 2025), and forms topologically associating domains (TADs) in the G1 nucleus (Schwarzer et al., 2017; Wutz et al., 2017). Although it is formally possible that local chromatin folding by cohesin contributes to the establishment of global CTs in early G1, a previous study has reported that cohesin depletion does not impair CT formation following endomitosis in human cells (Cremer et al., 2020). Whether and how condensin II and cohesin may functionally interplay to establish G1 CTs remains unknown.

**Figure 1.**
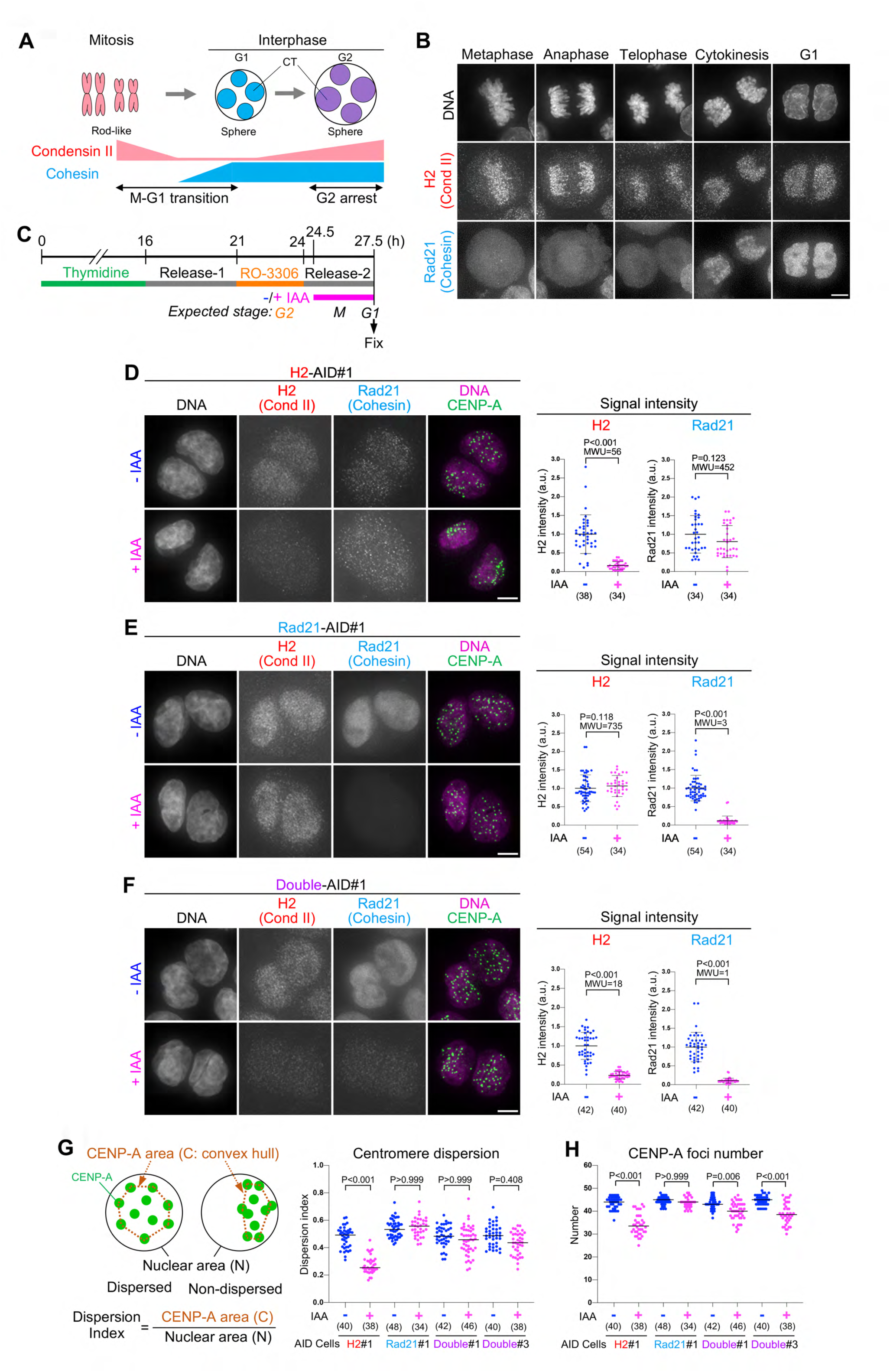
Condensin II depletion, but not cohesin or double depletion, suppresses post-mitotic centromere dispersion. **(A)** Conceptual outline of the current study. From late mitosis to early G1 phase, rod-shaped mitotic chromosomes undergo a morphological transition to spherical chromosome territories (CTs). During this transition, condensin II gradually dissociates from chromosomes and is progressively replaced by cohesin. In G2 phase, both condensin II and cohesin reside in the nucleus. This study focuses on two key stages, the M-to-G1 transition and G2 phase. (**B**) Localization of condensin II and cohesin during the M-to-G1 transition. Rad21-AID#1 cells expressing Rad21-mClover were cultured asynchronously, fixed, and immunolabeled with an anti-CAP-H2 antibody. DNA was counterstained with DAPI. Bar, 5 μm. **(C)** Schematic diagram of the cell preparation protocol for enriching G1 phase cells depleted of condensin II and/or cohesin. In brief, cells were first treated with thymidine, released, and subsequently treated with RO-3306 to arrest them at G2 phase. Following washout of RO-3306, cells progressed through mitosis into early G1, during which IAA (or no IAA) was added to deplete the target subunits. Enrichment of G1 cells using this protocol was assessed by FACS (Fig. S1 A). From the synchronized populations, early G1cells were selected based on their characteristic morphologies (i.e., pairs of small post-mitotic cells) and subjected to downstream analyses. Based on the measured nuclear sizes (Fig. S2 G), we confirmed that early G1 cells were appropriately selected. **(D-F)** (Left) H2-AID#1 (D), Rad21-AID#1 (E), and Double-AID#1 (F) cells were treated according to the protocol described in (C), fixed, and extracted as (B). H2-AID#1 cells were immunolabeled with antibodies against CAP-H2, Rad21, and CENP-A. Rad21-AID#1 cells and Double-AID#1 cells were immunolabeled with an anti-CAP-H2 antibody (Table S4). DNA was counterstained with DAPI. Bar, 5 μm. (Right) Nuclear signal intensities of CAP-H2 and Rad21 were measured and plotted. The numbers in parentheses at the bottom of each panel indicate the number of cells analyzed. P, p-value. MWU, the Mann-Whitney U test statistic. Bars represent the mean ± sd values. a.u., arbitrary unit. Data from an alternative Double-AID cell line (Double-AID#3) are shown in Fig. S2 F. **(G)** Quantification of CENP-A dispersion. (Left) Two representative patterns of CENP-A distribution (dispersed and non-dispersed) are illustrated. The centromere “dispersion index” is defined as the area of the convex hull encompassing the CENP-A signals divided by the nuclear area, as determined by DAPI staining. (Right) Dot plots show the centromere dispersion indices measured for the four AID cell lines examined. The numbers in parentheses at the bottom of the plot indicate the number of cells analyzed. **(H)** Dot plots show the number of CENP-A foci per nucleus. The numbers in parentheses at the bottom of the panel indicate the number of cells analyzed. Bars represent median values (G and H). The same set of images was used for quantifications shown in panels (G) and (H).

Does condensin II, which remains in the interphase nucleus, play an architectural role in maintaining interphase chromosome organization? In *Drosophila melanogaster,* condensin II restricts interchromosomal interactions (Hartl et al., 2008; Bauer et al., 2012), and contributes to the formation of spherical CTs through large-scale folding of interphase chromosomes (Hartl et al., 2008; Bauer et al., 2012; Rosin et al., 2018; Isenhart et al., 2025). In mammalian cells, it has been shown that condensin II associates with chromatin during S phase to initiate the resolution of newly duplicated sister chromatids in HeLa cells (Ono et al., 2013), and it also prevents hyper-clustering of pericentric heterochromatin and nucleoli in neural stem cells in mice (Nishide and Hirano, 2014). Despite these findings, there is currently no evidence that either condensin II, cohesin, or their combined action contributes to the maintenance of CT morphology in mammalian interphase cells (Cremer et al., 2020).

In the current study, we applied multiscale fluorescence in situ hybridization (FISH) techniques to a panel of human auxin-inducible degron (AID) cell lines that enable single or double depletion of condensin II and cohesin. Our analyses focused on two specific stages of the cell cycle, the M-to-G1 transition and G2 phase. During the M-to-G1 transition, condensin II depletion, but not cohesin depletion, caused defects in post-mitotic CT establishment. However, double depletion of condensin II and cohesin resulted in much more severe defects in the morphology of G1 CTs. During G2 phase, condensin II and cohesin were found to cooperate to maintain CT structures, although they act at different genomic scales. Remarkably, double depletion of both complexes led to severely deformed CTs that collapsed into the nucleolar periphery. Based on these findings, we discuss how the functional interplay between condensin II and cohesin contributes to the establishment and maintenance of large-scale chromosome architecture during interphase.

## Results

### Condensin II depletion, but not cohesin or double depletion, suppresses post-mitotic centromere dispersion

Previous studies established two auxin-inducible degron (AID) cell lines, CAP-H2-mACh and Rad21-mACl, in which CAP-H2 (a condensin II subunit) and Rad21 (a cohesin subunit) can be selectively depleted upon the addition of indole-3-acetic acid (IAA)(Natsume et al., 2016; Takagi et al., 2018)(Tables S1). In the current study, we referred to them as H2-AID#1 and Rad21-AID#1, respectively. To investigate how condensin II and cohesin contribute to chromosome structure, we generated a new cell line, CAP-H2-mACh/Rad21-mACl (hereafter Double-AID#1; Tables S2 and S3), in which condensin II and cohesin can be co-depleted. Moreover, to analyze centromere localization, we established three additional AID cell lines (H2-AID#2, Rad21-AID#2, and Double-AID#2), in which CENP-A-mClover is constitutively expressed (Tables S2 and S3).

We first examined the behavior of CENP-A signals in H2-AID#1, Rad21-AID#1, and Double-AID#1 cells during the M-to-G1 transition. In brief, cells were treated with thymidine, released, and subsequently arrested in G2 phase using RO-3306. After washing out RO-3306, the cells were allowed to progress from mitosis to early G1 phase for 3 hr in the presence or absence of IAA (Figs. 1 C and S1 A). The cells were then fixed and processed for immunofluorescence using antibodies against CENP-A to assess its nuclear distribution, and against CAP-H2 and Rad21 to evaluate the efficiency of protein depletion. Under control (-IAA) conditions in each cell line, CENP-A signals appeared as discrete foci that were evenly distributed throughout the G1 nucleus (Figs. 1, D-F and S2 F). We found that condensin II depletion suppressed post-mitotic centromere dispersion (Fig. 1 D), consistent with previous reports (Hoencamp et al., 2021; Sakamoto et al., 2022). In contrast, condensin I depletion did not suppress centromere dispersion (Fig. S2 A), suggesting that this suppression of post-mitotic centromere dispersion is not simply a secondary consequence of defects in chromosome assembly or segregation. Rather, it appears to be specifically caused by the absence of condensin II during the M-to-G1 transition. Live imaging further confirmed that centromere dispersion occurred during cytokinesis in control cells, but was suppressed in cells depleted of condensin II during the M-to-G1 transition (Fig. S2, C-E).

Apparently, little or no suppression of centromere dispersion was observed in cells depleted of cohesin alone or co-depleted of condensin II and cohesin (Figs. 1, E-F and S2 F).

To more precisely compare the degree of centromere dispersion under the three depletion conditions, we quantified two parameters. First, we measured the “dispersion index”, defined as the area of the convex hull encompassing the CENP-A foci divided by the nuclear area (Fig. 1 G). Second, we counted the number of distinguishable CENP-A foci, which tend to be high under dispersed conditions and lower when dispersion is suppressed (Fig. 1 H). The results showed that both parameters significantly decreased in condensin II-depleted cells, but not in cohesin-depleted cells. Interestingly, centromere dispersion was partially restored when cohesin was co-depleted with condensin II.

### Condensin II depletion, but not cohesin or double depletion, causes lengthwise elongation of G1 chromosomes

Based on the suppressed centromere dispersion observed in fixed cells (Fig. 1 D) and the chromosome morphology observed by live imaging (Fig. S2 D), we reasoned that condensin II depletion may cause chromosomes to retain a mitotic-like conformation without undergoing proper lengthwise shortening in post-mitotic cells, as previously suggested by computer simulations (Hoencamp et al., 2021). To test this possibility experimentally, we employed site-specific fluorescence in situ hybridization (FISH) probes targeting the centromere of chromosome 12 (Cen 12) and the 12q15 locus, which are ∼33 Mb apart (Fig. 2 A). The same set of cell lines (H2-AID#2, Rad21-AID#2, and Double-AID#2) was treated according to the protocol shown in Fig. 1 C, and subjected to FISH labeling using these site-specific probes. The distance between the two site-specific signals indeed increased upon condensin II depletion (Fig. 2, B and E), suggesting that condensin II contributes to the lengthwise compaction of chromosomes during the M-to-G1 transition. In contrast, cohesin depletion had a smaller effect on the distance between the two site-specific probes compared to condensin II depletion (Fig. 2, C and E). Double depletion did not result in a significant change, consistent with the partial restoration of centromere dispersion (Fig. 2, D and E).

**Figure 2.**
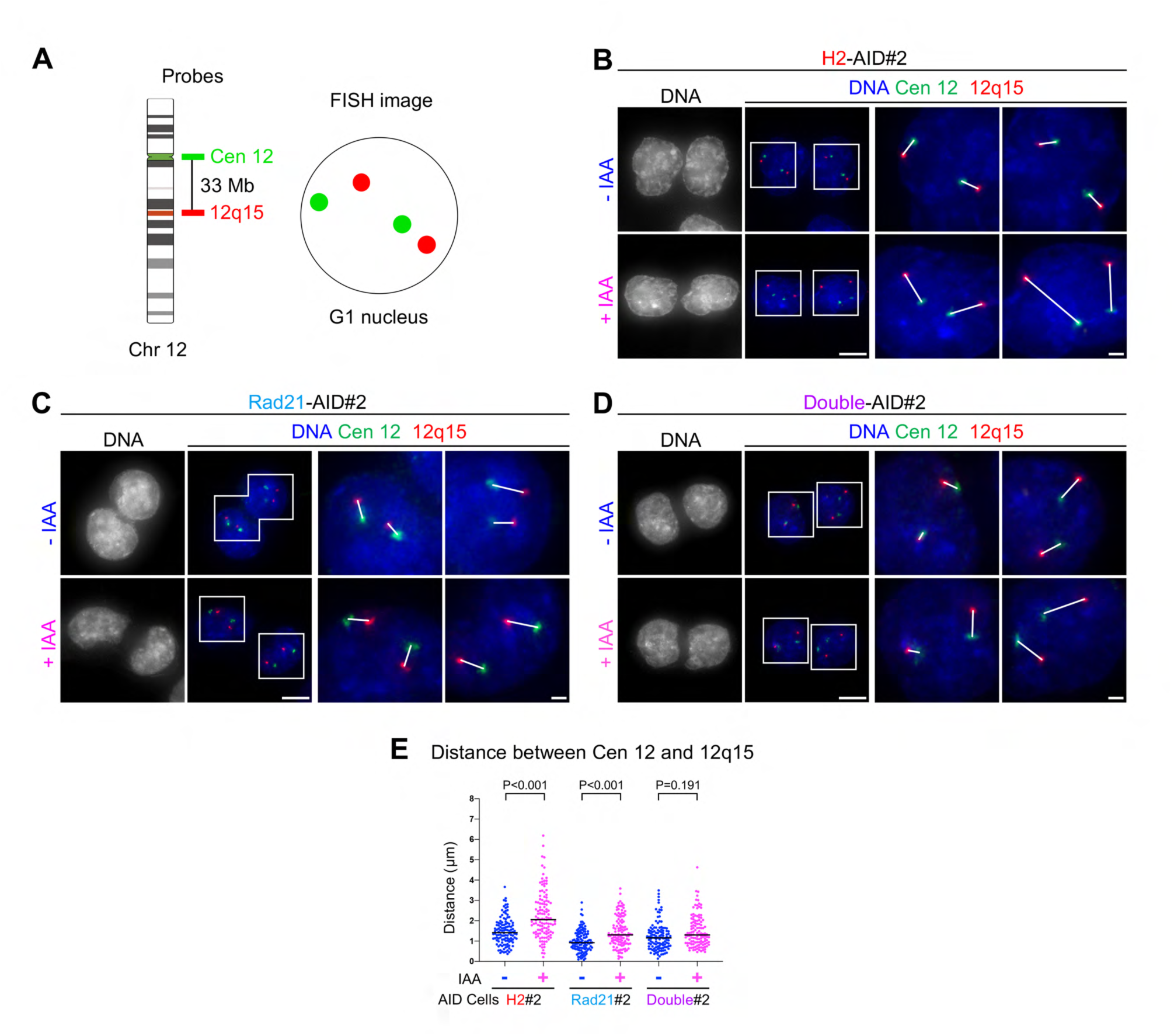
Condensin II depletion, but not cohesin or double depletion, causes lengthwise elongation of G1 chromosomes. **(A)** FISH probes used and a schematic of FISH signals expected. A mixture of a centromere 12 probe (Cen 12) and a site-specific probe for 12q15 was used (Table S5). The genomic distance between these two sites is 33 Mb. In diploid interphase cells, two pairs of Cen 12 (green) and 12q15 (red) signals are separately observed. **(B-D)** H2-AID#2 (B), Rad21-AID#2 (C), and Double-AID#2 (D) cells were treated as described in Fig. 1 C, fixed, and processed for FISH. DNA was counterstained with DAPI. Bar in the left panels, 5 μm. The regions enclosed by the white rectangles are magnified and shown on the right. Pairs of centromere 12 and 12q15 signals judged to be on the same chromosome are connected by white lines. Bar in right panel, 1 μm. **(E)** Dot plots show the distances between centromere 12 and 12q15 in three cell lines treated with (+) or without (-) IAA, as shown in panels (B-D). Bars represent median values. In this experiment, 120 signal pairs from 60 cells were analyzed under each condition.

### Double depletion of condensin II and cohesin causes severe morphological defects in G1 chromosome territories

During the M-to-G1 transition, individual chromosomes undergo a morphological transformation into roughly spherical structures, forming chromosome territories (CTs) within the reassembling nucleus (Cremer and Cremer, 2010). The above-mentioned findings prompted us to directly visualize CTs in cells depleted of condensin II, cohesin, or both in early G1. To this end, we performed FISH using whole chromosome painting (WCP) probes (Fig. 3 A, left). In human cells, chromosomes 18 and 19 exhibit distinct positioning within the nucleus (Tanabe et al., 2002; Cremer et al., 2003): homologs of chromosome 18 are typically separated and located near the nuclear periphery, whereas those of chromosome 19 tend to be closely associated and positioned in the nuclear interior (Fig. 3 A, right). We therefore reasoned that simultaneous FISH labeling of these two chromosomes would facilitate the detection of changes in their CT formation. Three cell lines (H2-AID#2, Rad21-AID#2, and Double-AID#2) were treated according to the protocol shown in Fig. 1 C, and subjected to FISH labeling. Under control conditions (-IAA), chromosomes 18 and 19 in each cell line exhibited spherical shapes and were distinctly localized within the nucleus, as expected (Fig. 3, B-D). Upon condensin II depletion, both chromosomes 18 and 19 adopted elongated, rod-shaped structures (Fig. 3 B [+IAA]). In contrast, no such characteristic structural changes were observed in cohesin-depleted cells (Fig. 3 C [+IAA]). Similar results were obtained using WCP probes for chromosomes 12 and 15 (Fig. S3, A-B and E-F). Notably, double depletion of condensin II and cohesin caused very severe defects in CT morphology across all examined chromosomes, resulting in expanded, cloud-like shapes (Figs. 3 D [+IAA] and S3, C and G [+IAA]). We speculate that the isotropic expansion of CTs in co-depleted cells may have obscured or masked the characteristic suppression of centromere dispersion (Fig. 1) and lengthwise elongation (Fig. 2) observed in cells depleted of condensin II alone.

**Figure 3.**
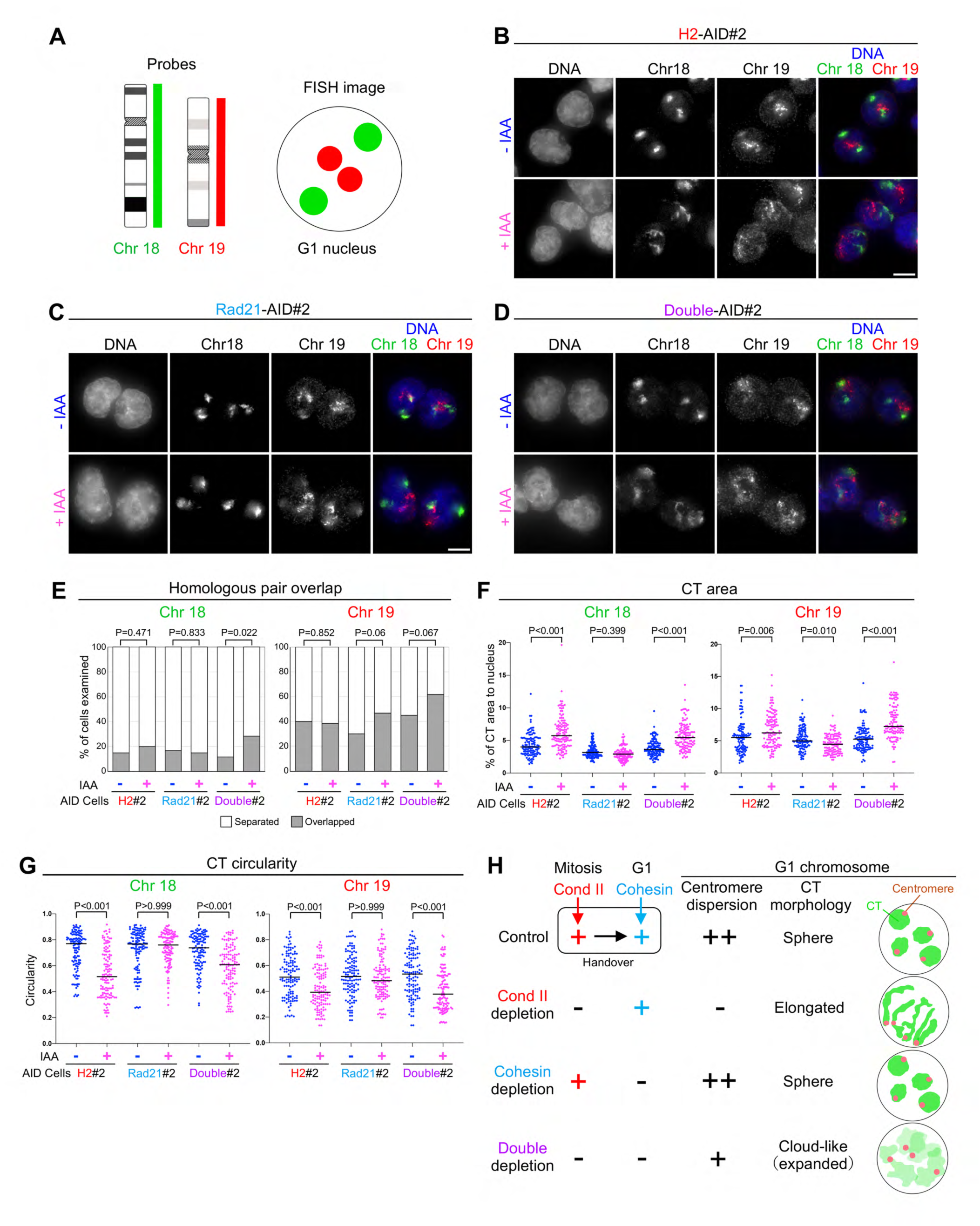
Double depletion of condensin II and cohesin causes severe morphological defects in G1 chromosome territories. **(A)** FISH probes used and a schematic of FISH signals expected. Whole chromosome painting (WCP) probes for human chromosomes 18 and 19 were used. The schematic diagram of human chromosomes was adapted with minor modifications from the instruction manual provided by the manufacturer of the FISH probes (MetaSystem probes, Altlussheim, Germany). **(B-D)** H2-AID#2 (B), Rad21-AID#2 (C), and Double-AID#2 (D) cells were treated according to the protocol described in Fig. 1 C, fixed, and processed for WCP FISH as described in Materials and Methods. DNA was counterstained with DAPI. Bar, 5 μm. **(E)** Bar graphs show the proportion of cells with overlap between chromosomes 18 homologs and 19 homologs. **(F)** Dot plots show the area of FISH-labeled territories for chromosomes 18 and 19. **(G)** Dot plots show the circularity of FISH-labeled territories for chromosomes 18 and 19. Bars represent median values (F and G). The same set of images was used for quantifications in (E), (F) and (G), with 120 chromosomes (from 60 cells) in each condition. (H) Summary of the key observations during the M-to-G1 transition.

We then quantified the changes in CT morphology by measuring three parameters: the overlap between homologous pairs, CT area, and CT circularity. Circularity is a morphological index that reflects how closely the shape of an object approximates a perfect circle, with a value of 1 indicating a perfect circle. The data showed that double depletion increased the overlap between homologs of both chromosomes 18 and 19 (Fig. 3 E), consistent with the observed expansion of their CTs. Chromosome 19 homologs exhibited a higher probability of overlap even in control cells, likely reflecting their characteristic positioning in the nuclear interior. Moreover, we found that CT area increased (Figs. 3 F and S3, D and H) and CT circularity decreased (Fig. 3 G) upon condensin II depletion and double depletion, but not upon cohesin depletion. Based on these results, we speculate that centromere dispersion during the M-to-G1 progression (Fig. 1) is mechanistically linked to the structural conversion of rod-shaped chromosomes into spherical CTs (Figs. 2 and 3), both of which are impaired in condensin II-depleted cells. The apparent recovery of centromere dispersion observed upon co-depletion with cohesin (Fig. 1 F) is likely a secondary effect, attributable to the severe defects in CT formation observed under this condition (Fig. 3 D).

The key observations during the M-to-G1 transition are summarized in Fig. 3 H. Condensin II facilitates the morphological transition of chromosomes from rod-like to spherical shapes, thereby promoting the establishment of proper CTs in early G1.

While cohesin depletion alone has little effect on CT formation, double depletion of condensin II and cohesin results in severe defects in CT morphology, distinct from those caused by condensin II depletion alone. We propose that the sequential actions of condensin II and cohesin, constituting a functional “handover”, are crucial for the proper establishment of post-mitotic CTs.

### Cohesin depletion, but not condensin II depletion, causes defects in local chromatin folding in G2-arrested cells

Next, we sought to elucidate the functional contributions of condensin II and cohesin to chromosome organization during G2 phase when both complexes are present in the nucleus. In this section, we focus on chromatin organization at the ∼1 Mb scale, hereafter referred to as “local chromatin folding”. We employed a pair of site-specific FISH probes targeting at 11q13.3, positioned 0.56 Mb apart (Fig. 4 A; P, proximal; D, distal). Because G2 cells have duplicated sister chromatids, the two probes have a potential to detect a pair of P signals (red) and a pair of D signals (green). However, sister signals were distinguishable only under cohesin and double depletion conditions (Fig. 4, D and E). In the following discussion, we focus on the distance between P and D signals to evaluate local chromatin folding. The degree of sister chromatid separation (i.e., P-P and D-D distances) under each condition is presented in Fig. S5, A and B. See the legend of Fig. 4 A for details on how separated sister FISH signals were handled in the quantitative analysis described below.

**Figure 4.**
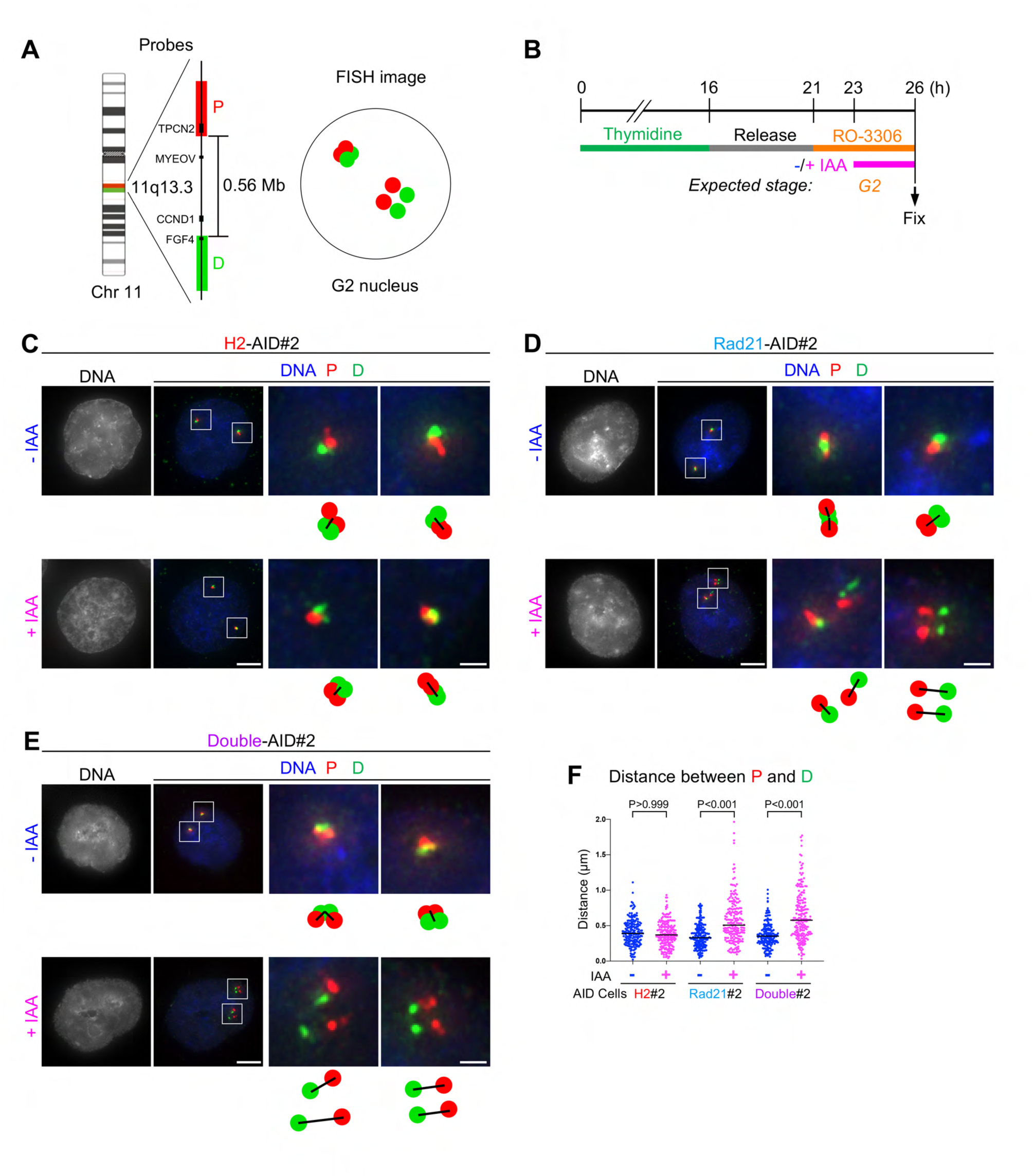
Cohesin depletion, but not condensin II depletion, causes defects in local chromatin folding in G2-arrested cells. **(A)** FISH probes used and a schematic of FISH signals expected. A pair of site-specific probes targeting 11q13.3 were used. The genomic distance between the two probes was 0.56 Mb. The cytological distances between P and D were measured, yielding two values per chromosome in G2 cells (e.g., the right signal set shown in the FISH image cartoon). When sister signals could not be distinguished, the same distance was assigned to two chromatids (e.g., the left signal set; see also H2-AID#2 panels [C] or control [-IAA] panels). In cohesin and double depletion, sister signals were frequently observed as separated. The distances between the P signals and between the D signals (i.e., the degree of sister chromatid resolution) are shown in Fig. S5. **(B)** Schematic diagram of the cell preparation protocol for G2-arrested cells depleted of condensin II and/or cohesin subunits. In brief, cells were first treated with thymidine, released, and subsequently treated with RO-3306 to arrest them at G2 phase. Two hours after RO-3306 addition, IAA (or no IAA) was added to deplete the target subunits. Enrichment of G2 cells using this protocol was assessed by FACS analysis (Fig. S1 B). **(C-E)** H2-AID#2 (C), Rad21-AID#2 (D), and Double-AID#2 (E) cells were treated according to the protocol described in (C), fixed, and processed for FISH using a pair of site-specific probes targeting 11q13.3. DNA was counterstained with DAPI. Bar in the left panels, 5 μm. The regions enclosed by the white rectangles are magnified and shown on the right. Pairs of P (red) and D (green) judged to be on the same chromosome are connected by black lines in the cartoons shown at the bottom. Bar in right panel, 1 μm. **(F)** Dot plots show the distances between P and D. Bars represent median values. In this experiment, 200 signal pairs from 50 cells were analyzed under each condition.

Three cell lines (H2-AID#2, Rad21-AID#2, and Double-AID#2) were synchronized in G2 phase using thymidine and RO-3306 (Figs. 4 B and S1 B), followed by treatment with or without IAA for 3 hr (Fig. S4, A-D). We found that, under control conditions, the P and D signals overlapped or were located in close proximity (Fig. 4, C-E [-IAA]). A similar distribution of the signals was observed in condensin II-depleted cells (Fig. 4 C [+IAA]). In contrast, the distance between P and D signals significantly increased in cohesin-and co-depleted cells (Fig. 4 D [+IAA], and E [+IAA]), suggesting that local chromatin folding was compromised under these conditions (Fig. 4 F). We interpreted these cytological observations as reflecting the destabilization of TADs caused by cohesin depletion, as previously revealed by Hi-C and FISH analyses (Rao et al., 2017; Luppino et al., 2020). Importantly, such alternations of local chromatin folding were not observed upon condensin II depletion. We also tested additional pairs of commercially available probes and obtained similar results (Fig. S5, C-F).

### Condensin II depletion, but not cohesin depletion, causes defects in 20 Mb-scale chromosome arm conformation in G2-arrested cells

Next, we sought to examine G2 chromosome arm conformation under conditions of condensin II and/or cohesin depletion. To this end, we prepared a set of three FISH probes targeting the short arm of chromosome 17: a mixture of centromere-specific conventional probes (C) and two custom site-specific oligo probes (P and D3)(Fig. 5 A and Table S6). The mixture of these probes simultaneously detects the centromeric region of chromosome 17 (C, yellow), a proximal site located 5 Mb from the centromere (P, green), and a distal site located 20 Mb from the centromere (D3, red)(see Table S7 for details). To assess the conformation of the chromosome 17 short arm, we analyzed the angle formed at the P site (“P angle”; Fig. 5 B). The P angle value reflects the degree of bending of the chromosome arm within this 20 Mb region.

**Figure 5.**
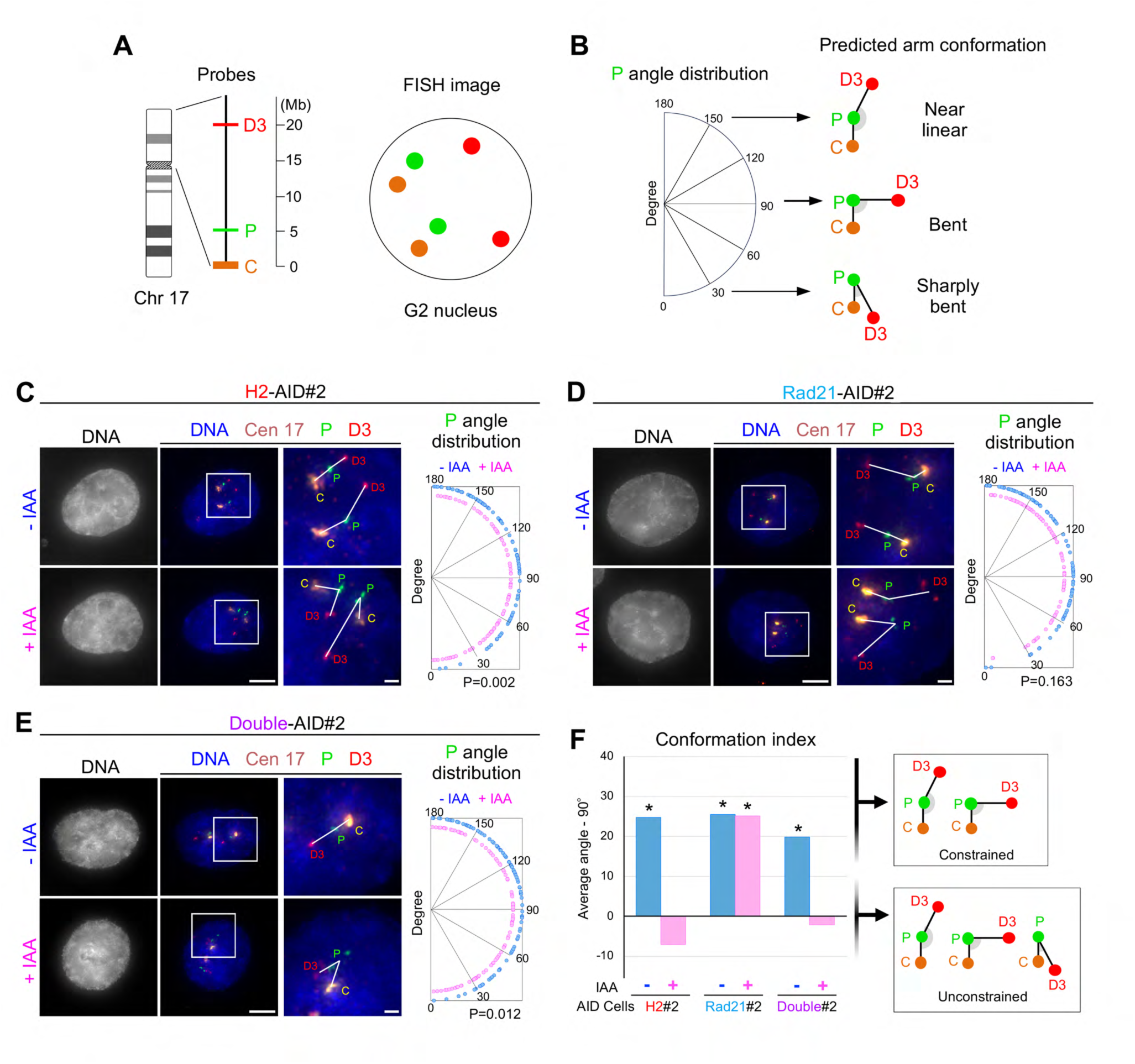
Condensin II depletion, but not cohesin depletion, causes defects in 20 Mb-scale chromosome arm conformation in G2-arrested cells. **(A)** FISH probes used and a schematic of FISH signals expected. In this analysis, a set of three site-specific probes was used: two custom oligo-probes targeting specific loci (P and D3) on the short arm of chromosome 17, and a mixture of chromosome 17 centromere-specific probes (Table S7). Oligo probes P and D3 locate at 5 Mb and 20 Mb away from the centromere, respectively. Consequently, two groups of FISH signals including centromeres (yellow), P sites (green), and D3 sites (red) are delineated within a diploid nucleus, as shown in the right panel. (**B**) Assessment of chromosome arm conformation based on the P angle. The P angle is defined as the angle formed between the lines connecting P to D3 and P to C. It reflects the degree of bending of the chromosome arm in this region. For example, a P angle greater than 120° indicates a nearly linear arm, an angle of 90° indicates moderate bending, and an angle of 30° indicates a sharply bent arm. (**C-E**) H2-AID#2 (C), Rad21-AID#2 (D), and Double-AID#2 (E) cells were treated as described in Fig. 4, C-E, fixed, and processed for FISH. DNA was counterstained with DAPI. Bar in the left panels, 5 μm. The regions enclosed by the white rectangles are magnified and shown on the right. A set of C (yellow), P (green) and D3 (red) judged to be on the same chromosome is connected by white lines. The position of each signal was defined as the centroid of the corresponding FISH-positive area. When sister FISH signals were separated, an average value of their centroid coordinates was used for analysis. Bar in right panel, 1 μm. P angle distributions in the cells treated with (+) or without (-) IAA are shown as circular scatter plots (right). Statistical significance was assessed using the Mardia Watson Wheeler test. In this experiment, 100 signal sets (i.e. chromosomes) from 50 cells were analyzed under each condition. **(F)** Quantification of the P angle distribution. The bar graphs show the “conformation index”, defined as the average angle P minus 90° (See text). Deviations from randomness were assessed using the Rayleigh test. Asterisks indicate statistically significant deviations from a random distribution (P< 0.05). In this analysis, 100 sets of signals (from 50 cells) were examined under each condition. The data from a representative experiment set out of three repeats are shown.

Three cell lines (H2-AID#2, Rad21-AID#2, and Double-AID#2) were treated using the same protocol as described in Fig. 4 A, and subjected to FISH labeling. Under control conditions, although the P angle values were distributed across a wide range, chromosomes with sharp bends (i.e., angle values below 30°) were rarely observed (Fig. 5, C-E [-IAA]). In contrast, the frequency of sharply bent chromosomes significantly increased upon condensin II depletion (Fig. 5 C [+IAA]), but not upon cohesin depletion (Fig. 5 D [+IAA]). Similar to condensin II depletion, double depletion also resulted in an increased frequency of sharply bent chromosomes (Fig. 5 E [+IAA]). Fig. 5 F shows a plot of the “conformation index”, defined as the average angle P minus 90°, which indicates the extent to which the P angle distribution deviates from a random distribution. This index was high under control conditions, where sharply bent chromosomes were rare, but decreased markedly under condensin II and double depletion conditions, where sharply bent chromosomes became more frequent. These results suggest that condensin II imposes structural constraints on chromosome arm conformation at the ∼20 Mb scale examined.

We then asked to what extent condensin II-dependent arm conformation observed at the ∼20 Mb scale could also apply at different genomic scales. To address this, we performed the same analysis using additional sets of FISH probes spanning different genomic intervals (Fig. S6). We found that the conformation index was close to zero or below zero at scales below 15 Mb or above 40 Mb scale, and was not significantly affected by condensin II depletion (Fig. S6 I). Thus, these results suggest that condensin II plays a crucial role in determining chromosome arm conformation at a genomic span of ∼20 Mb, a scale larger than that typically regulated by cohesin (Fig. 4). Physiological implications of this finding are discussed in the Discussion section below.

### Double depletion of condensin II and cohesin results in abnormal perinucleolar distribution of CTs in G2-arrested cells

Next, we tested whether depletion of condensin II and/or cohesin affects the maintenance of CT morphology in G2-arrested cells, using WCP FISH probes targeting multiple chromosomes. In these experiments, we employed the same synchronization protocol as described above, but extended the duration of IAA treatment from 3 to 5 hr (Fig. 6 A). The efficiencies of G2 phase synchronization and target protein depletion were comparable between the 3 hr and 5 hr IAA treatments (Figs. S1 B and S7). We found that condensin II or cohesin depletion alone had little if any effect on CT morphology. However, double depletion resulted in severe morphological defects in the CTs of chromosomes, 18, 19, 21, and 22, frequently producing irregular crescent shapes (Fig. S8, C-D). Remarkably, these abnormal CTs were frequently displaced toward the nucleolar periphery, as judged by DAPI staining.

**Figure 6.**
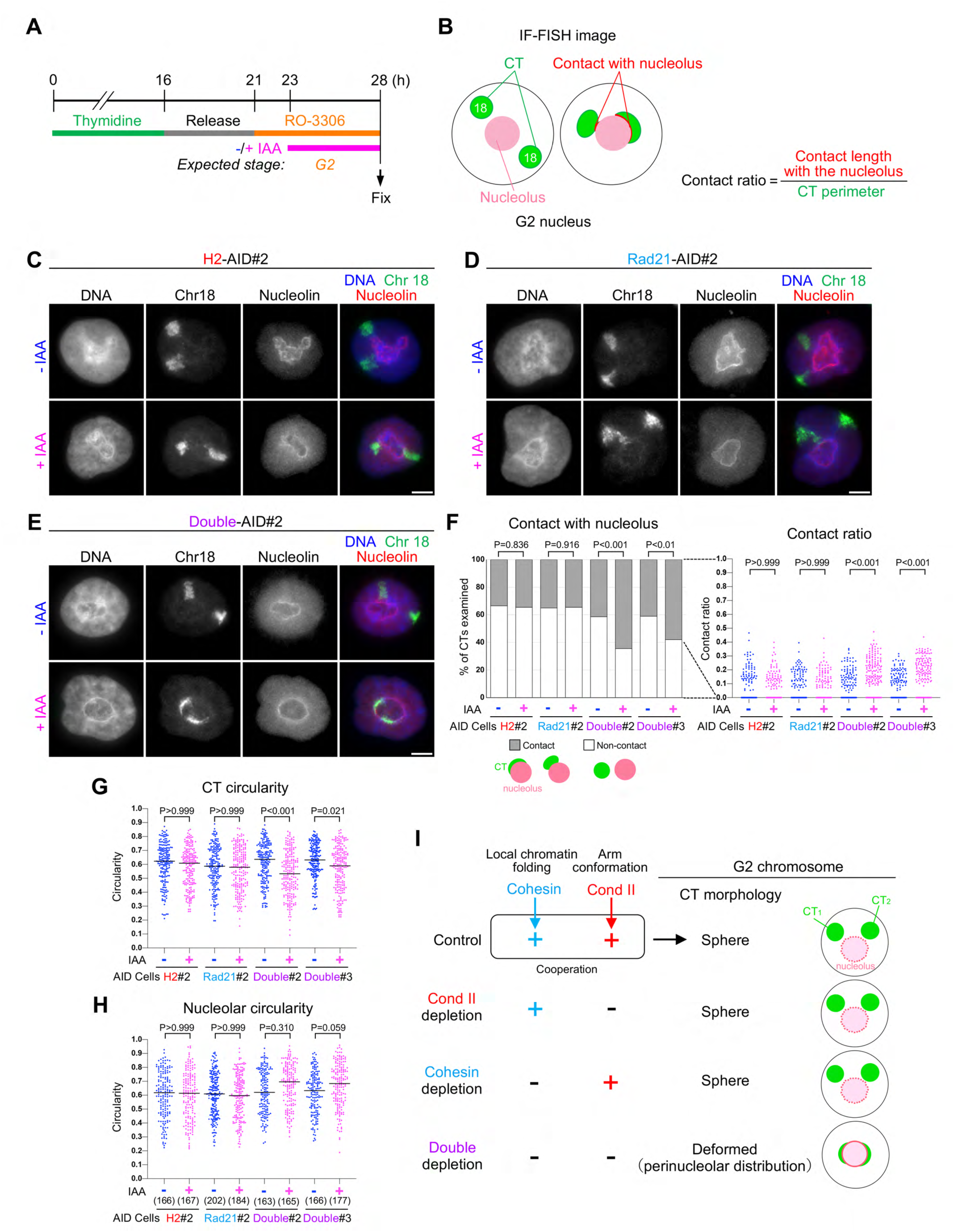
Double depletion of condensin II and cohesin results in abnormal perinucleolar distribution of CTs in G2-arrested cells. **(A)** Schematic diagram of the cell preparation protocol for analyzing CT morphology in G2 arrested cells co-depleted of condensin II and cohesin subunits. The same synchronization protocol described in Fig. 4 B was used, except that the duration of IAA treatment was extended from 3 hr to 5 hr. The extent of G2 arrest was confirmed by FACS analysis (Fig. S1 B). **(B)** IF-FISH image and quantification method for deformed CTs around the nucleolus. In this experiment, the nucleolin-positive area was defined as the nucleolus. Under unperturbed condition, the chromosome 18 homologs were separated and located near the nuclear periphery, without associating with nucleoli (left panel). In cells co-depleted of condensin II and cohesin, a characteristic perinucleolar distribution of the chromosome 18 homologs was observed. To assess the extent of surface contact between chromosome 18 and nucleoli, we defined the CT-nucleolus contact ratio as the length of the CT perimeter in contact with nucleoli divided by the total perimeter. **(C-E)** H2-AID#2 (C), Rad21-AID#2 (D), and Double-AID#2 (E) cells were treated as described in (A), and fixed according to the procedure in Fig. 2, B-D. After extraction, cells were immunolabeled with an anti-nucleolin antibody, re-fixed with 2 % PFA in PBS, and subsequently hybridized with WCP probes for chromosomes 18 (Tables S4 and S5). Bar, 5 μm. **(F)** Quantification of chromosome 18-nucleolus contact. Bar graphs (left) show the proportion of CTs with surface contact between chromosome 18 and nucleoli. Dot plots (right) show the distribution of the contact ratio, as defined in (B). Bars represent median values. **(G)** Quantification of CT morphology. Dot plots show the circularity of the FISH signals for chromosomes 18 in the four AID cell lines. The same images were used for quantifications in (F) and (G), with 200 CTs from 100 cells under each condition. IF-FISH images from an alternative Double-AID cell line (Double-AID#3) are shown in Fig. S8 A. **(H)** Quantification of nucleolar morphology. Dot plots show the circularity of the nucleolin-positive area in the four AID cell lines. The number of cells analyzed for nucleolar morphology was the same as in (G); however, because the number of nucleoli varies between nuclei, the actual number of nucleoli examined is indicated in parentheses. Bars represent median values. **(I)** Summary of the key observations during G2 phase.

To validate the relative positioning of the abnormal CTs and the nucleolus more precisely, we performed immunofluorescence-FISH (IF-FISH) using a combination of an anti-nucleolin antibody and a WCP probe for chromosome 18 (Fig. 6 B, left). Under control conditions, ∼30% of chromosome 18 territories exhibited partial contact with the nucleolus (Fig. 6, C-F [-IAA]). Although depletion of either condensin II or cohesin alone had little effect on the contact frequency (Fig. 6, C, D and F [+IAA]), double depletion of condensin II and cohesin markedly increased it in two independent cell lines (Double-AID#2 and Double-AID#3) tested (Figs. 6, E and F [+IAA], and S8 A, [+IAA]). Under the same conditions, no overlap between chromosome 18 homologs was observed (Fig. S8 B). To further evaluate the CT-nucleolus contacts, we quantified the “contact ratio”, defined as the length of the CT perimeter in contact with the nucleolus divided by the total CT perimeter (Fig. 6 B, right), and found that this parameter was significantly increased under double depletion conditions (Fig. 6 F).

Finally, we assessed another morphological index, circularity, for both CTs and nucleoli. While condensin II or cohesin depletion alone did not affect the circularity of either structure, double depletion results in a decrease in CT circularity and an increase in nucleolar circularity (Fig. 6, G and H). These results further support the notion that CTs undergo substantial deformation in the absence of both condensin II and cohesin.

Curiously, the nucleoli displayed smoother surfaces under these conditions, likely due to the relaxation of physical constrains normally imposed by surrounding CTs.

The key observations during G2 phase are summarized in Fig. 6 I. Cohesin and condensin II contributes to G2 chromosome organization at distinct genomic scales: cohesin regulates local chromatin folding at the ∼1 Mb scale, whereas condensin II governs chromosome arm conformation at the ∼20 Mb scale. Although depletion of either cohesin or condensin II alone has little effects on large-scale CT morphology, double depletion results in severe morphological defects, including abnormal perinucleolar distribution of CTs.

## Discussion

Although it is well established that condensin II mediate lengthwise chromosome compaction during mitosis (Ono et al., 2003; Gibcus et al., 2018; Samejima et al., 2025), its potential role in interphase chromosome organization remains poorly defined. The initial motivation of the current study was to elucidate the functional contribution of condensin II to interphase chromosome organization, particularly in relation to cohesin. To our knowledge, this is the first study to systematically compare the effects of condensin II, cohesin, and double depletion in the same experimental system. By focusing on two distinct stages of the cell cycle: the mitosis-to-G1 transition (Figs. 1-3) and G2 phase (Figs. 4-6), we provide new insights into the dynamic interplay between these two SMC protein complexes.

### Functional handover from condensin II to cohesin contributes to the establishment of G1 chromosome territories

A previous study reported that condensin II depletion results in the retention of Rabl-like conformations, as judged by Hi-C in haploid (HAP1) cells, and centromere clustering, as observed by cytology in both Hap1 and diploid cells (Hoencamp et al., 2021). In the current study using human diploid cells, we confirm that condensin II depletion suppresses post-mitotic centromere dispersion (Fig. 1) and further demonstrate that this defect is accompanied by a failure of proper formation of G1 CTs (Figs. 2 and 3). Live-cell imaging further reveals that centromere dispersion is tightly linked to global chromosome reorganization during the M-to-G1 transition (Fig. S2 D). Together, these findings underscore an essential role of condensin II in the establishment of CTs after mitosis.

While cohesin depletion alone had little impact on centromere dispersion and CT formation in G1 nuclei, we were surprised to find that double depletion of condensin II and cohesin caused severe defects in CT establishment and an apparent restoration of centromere dispersion (Figs. 1-3). These phenotypes, distinct from those caused by condensin II depletion alone, suggest that cohesin plays a previously unrecognized role in CT establishment that becomes evident only when condensin II is compromised.

Condensin II-depleted chromosomes may not serve as optimal substrates for cohesin in early G1. We therefore propose that a functional “handover” from condensin II to cohesin is critical for the establishment of CTs in early G1 (Figs. 3 H and 7 A). By employing new cell lines that allow double depletion of condensin II and cohesin, our current study extends previous Hi-C (Abramo et al., 2019) and imaging (Brunner et al., 2025) analyses of the M-to-G1 transition. The important question of to what extent condensin II and cohesin “co-occupy” chromatin during this transition remains to be fully clarified.

### Differential contributions of condensin II and cohesin to the maintenance of G2 chromosome territories

To dissect the functional interplay between cohesin and condensin II in the maintenance of G2 chromosome organization, we established FISH-based assays at three different genomic scales: local chromatin folding (∼1 Mb scale), chromosome arm conformation (∼20 Mb scale) and whole CTs (>50 Mb scale).

First, local chromatin folding (∼1 Mb scale) was assessed by measuring the distance between two closely positioned loci labeled by FISH (Fig. 4). Our results using probes for 11q13.3 show that cohesin, but not condensin II, is essential for the maintenance of local chromatin folding. This finding perfectly complements previous Hi-C data showing that cohesin depletion causes a loss of TAD structure at these loci (Rao et al., 2017). The increased distance of the two FISH signals observed here likely reflect such TAD disorganization (Luppino et al., 2020). While earlier imaging studies examined cohesin-dependent chromatin folding at higher resolution (Bintu et al., 2018; Finn et al., 2019; Gabriele et al., 2022; Sun et al., 2023), our study is the first to directly compare the effects of condensin II depletion, cohesin depletion and double depletion in parallel. Although not the primary focus of the current study, our FISH-based assay also provides valuable information about the extent of sister chromatid resolution in the examined loci. Brief analyses of sister distances are presented in Fig. S5.

Second, chromosome arm conformation (∼20 Mb scale), was assessed by measuring the extent of arm bending using FISH probes targeting two specific loci and the centromere on chromosome 17 (Fig. 5). In control cells, the P angle on the short arm of chromosome 17 showed a non-random distribution. Condensin II depletion and double depletion, but not cohesin depletion, increased the frequency of sharply bent conformations, resulting in a randomized distribution of P angle. These findings suggest that condensin II, but not cohesin, constrains chromosome arm conformation at this genomic scale.

Third, whole CT morphology (>50 Mb scale) was assessed using WCP probes (Fig. 6). A previous study reported that, under the special conditions following endomitosis, cohesin depletion causes little or no alteration in CT morphology, even within a multilobulated nucleus (Cremer et al., 2020). Consistently, our current results show no discernible defects in CT morphology upon depletion of either cohesin or condensin II alone. Strikingly, however, double depletion of cohesin and condensin II lead to severe CT deformation, revealing previously underappreciated roles of these SMC protein complexes in large-scale G2 chromosome organization. Taken together, our findings demonstrate that although cohesin and condensin II act at distinct genomic scales, their cooperative functions are essential for preserving CT morphology during G2 phase (Figs. 6, 7, and S8).

**Figure 7.**
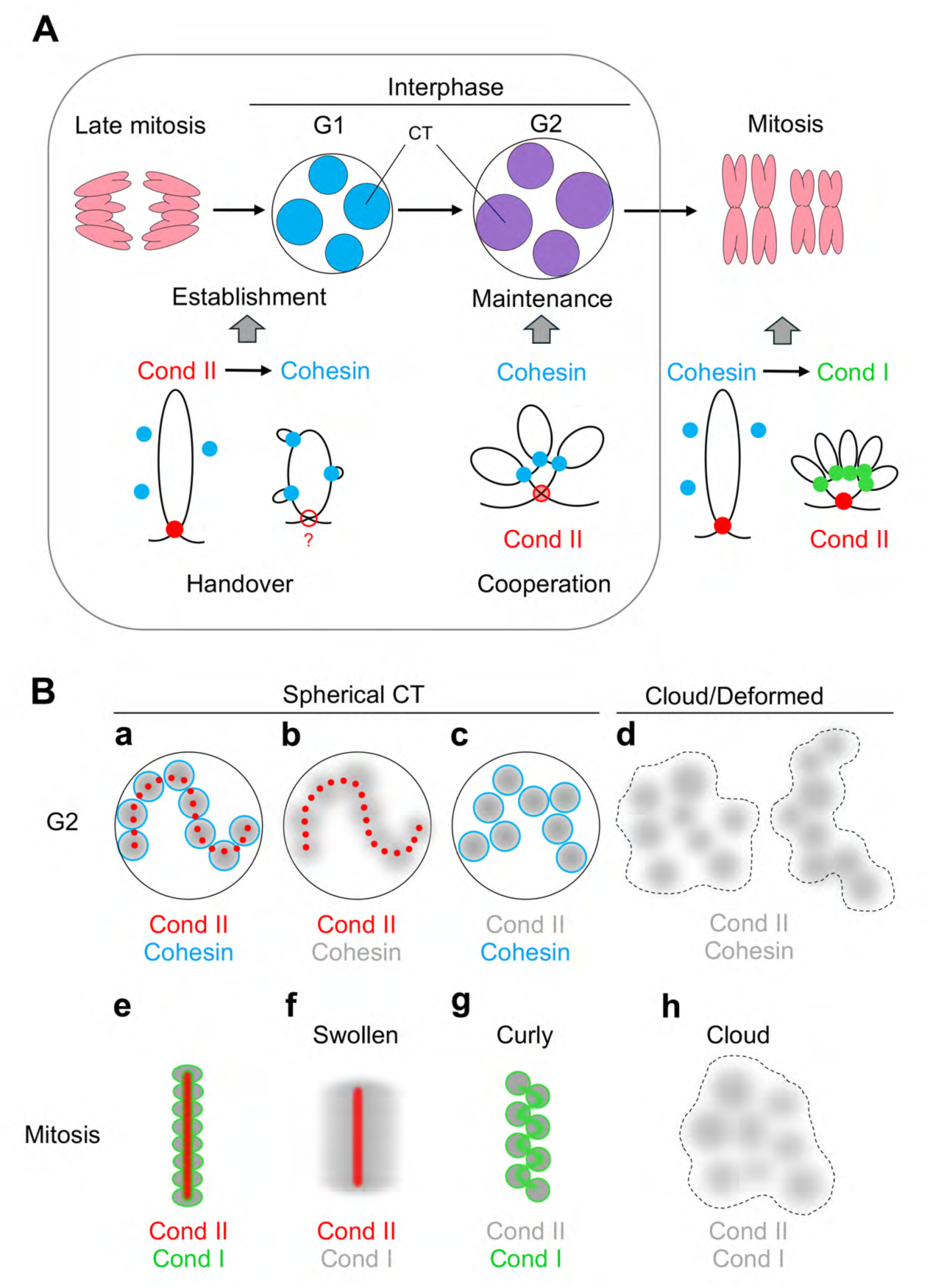
Models for the orderly and cooperative actions of SMC protein complexes. **(A)** Dynamic interplay between SMC protein complexes throughout the cell cycle. At the M-to-G1 transition, the contribution of condensin II to mitotic chromosome organization gradually diminishes, although its residual activity likely persist to support post-mitotic chromosome architecture. In early G1, cohesin begins to promote local chromatin folding, including the formation of TADs. This sequential action, referred to as a functional “handover” from condensin II to cohesin, is important for the establishment of interphase CTs. During G2 phase, condensin II and cohesin cooperate to maintain CT morphology, each acting at different genomic scales. At the G2-to-M transition, cohesin dissociates from chromatin, while mitotically activated condensin II initiates its mitosis-specific functions. This is followed by condensin I-mediated nested loop formation, thereby completing metaphase chromosome assembly. The key cell cycle stages addressed in the current study are indicated by the large square. **(B)** Comparison between G2 territories (a-d) and mitotic chromosomes (e-h). In G2 cells, depletion of cohesin (b) or condensin II (c) impairs local chromatin folding or chromosome arm conformation, respectively, but has little impact on overall CT morphology. These phenotypes may resemble the “swollen” and “curly” chromosomes observed in mitotic cells depleted of condensin I (f) and condensin II (g), respectively (Ono et al., 2003). In contrast, double depletion of cohesin and condensin II in G2 cells (d) results in severely deformed CTs, which resemble the “fuzzy” or cloud-like chromosomes observed in mitotic cells co-depleted of condensins I and II (h)(Ono et al., 2003). These observations suggest that the functional interplay between condensin II and cohesin in G2 phase may parallel that between condensin II and condensin I in mitosis.

What underlies the abnormal perinucleolar distribution of CTs observed under the double depletion condition? We speculate that double depletion of condensin II and cohesin in human cells markedly reduces chromatin constraints operating at distinct genomic scales, thereby allowing CTs to accumulate around nucleoli via depletion forces (Iida et al., 2024). Under the same conditions, nucleoli exhibit smoother surfaces, likely due to the release of physical constraints normally imposed by surrounding chromosomes (Fig. 6 E). Interestingly, the perinucleolar accumulation resembles the “surrounded nucleolus (SN)” type observed transiently in mouse oocytes and 2-cell stage embryos (Bonnet-Garnier et al., 2012; Borsos and Torres-Padilla, 2016; Nakaya et al., 2017). Although both condensin II and cohesin are present in oocyte nuclei (Lee et al., 2011), their nuclear volume extremely large, over 100 times that of human nuclei used in the current study (Wesley et al., 2020; Kyogoku and Kitajima, 2023), suggesting that a low-crowding nuclear environment may similarly promote SN-type CT formation through depletion forces. The notion that chromatin crowding strongly influences CTs may also explain previous observations that condensin II deficiency alone alters CT morphology in *Drosophila melanogaster* (Bauer et al., 2012; Rosin et al., 2018; Isenhart et al., 2025). The differential effects may be attributable to the small genome size and lower nuclear DNA density of *Drosophila* compared to humans (Hara et al., 2013).

Moreover, the functional contributions of cohesin and boundary factors to interphase genome architecture differ between *Drosophila* and humans (Rowley et al., 2017; Matthews and White, 2019), providing a potential explanation for the divergent observations.

### Dynamics of SMC protein complexes throughout the cell cycle

It is important to evaluate our current findings from a broader perspective that considers chromosomal events occurring across different stages of the cell cycle. For example, the functional handover from condensin II to cohesin during the M-to-G1 transition appears to be “reversed” during the G2-to-M transition (Gibcus et al., 2018; Samejima et al., 2025). In the latter transition, bulk cohesin dissociates from chromatin, while mitotically activated condensin II initiates mitotic chromosome assembly, a process that is subsequently completed by condensin I (Fig. 7 A). Comparative analysis of these two transitions may provide deeper insights into the dynamic reorganization of chromosomes across the cell cycle.

Our assay identified the ∼20 Mb scale as the most sensitive window for detecting condensin II-dependent maintenance of chromosome arm conformation (Figs. 5 and S6). This finding raises the intriguing possibility that the ∼20 Mb scale represents a fundamental unit of the structural backbone of chromosomes, whose integrity depends on condensin II. We speculate that this unit is inherited from mitotic chromosomes following their disassembly in late mitosis, and subsequently serves as a building block for chromosome axis reassembly in the next mitosis. A hallmark phenotype of condensin II depletion in mitosis is the formation of “zig-zag” or “curly” chromosomes, a defect that arises despite the presence of condensin I (Ono et al., 2003; Green et al., 2012). In contrast, double depletion of condensins I and II leads to a much more severe defect, resulting in amorphous, cloud-like chromatin masses (Ono et al., 2003). Notably, the functional interplay between condensin II and cohesin in G2 phase described here may parallel the interplay between condensin II and condensin I in mitosis (Fig. 7 B). If this is the case, condensin II may function as a central organizer of higher-order chromosome architecture throughout the cell cycle, with cohesin and condensin I imposing additional physical constraints on chromatin by forming DNA loops in an interphase-and mitosis-specific manner, respectively. Finally, it should be stressed that the “backbone” discussed above, while possibly involving condensin-condensin interactions (Kinoshita et al., 2022), is not a static or rigid structural element. Rather it represents a dynamic framework that may underlie the condensin II-dependent helical staircase (Gibcus et al., 2018) or the disordered, discontinuous helical organization (Samejima et al., 2025) inferred from Hi-C analyses.

In the current study, we have uncovered previously unrecognized aspects of the functional interplay between condensin II and cohesin. Although our analysis was limited to selected chromosomal regions, future genome-wide analyses will be essential to assess the generality of these findings. Our results complement and substantially extend the emerging concept that the orderly actions of SMC protein complexes orchestrate chromatin dynamics throughout the cell cycle. These include: (1) a functional handover from condensin II to cohesin for the establishment of G1 CTs (Abramo et al., 2019; Brunner et al., 2025)(this study); (2) the collaborative maintenance of G2 CTs by cohesin and condensin II (this study); and (3) a reverse handover from cohesin to condensins in early mitosis for chromosome assembly (Losada et al., 2002; Shintomi and Hirano, 2011; Gibcus et al., 2018; Samejima et al., 2025)(Fig. 7A). A future challenge is to elucidate how these sequential actions are regulated by diverse extrinsic factors and a multitude of post-translational modifications during the cell cycle and beyond.

## Materials and Methods

### Cell lines

The auxin-inducible degron (AID) cell lines used in this study are listed in Table S1. All AID cell lines used in this study were presumed to maintain diploidy (Fig. S1). Double-AID cell lines enabling double depletion of condensin II and cohesin were established using CRISPR/Cas9-mediated genome editing. Donor plasmids targeting the Rad21 locus (pMK265 and RAD21 C-tag CRISPR) were transfected into H2-AID#1 cells using FuGene HD (Promega, Madison, WI, USA)(Natsume et al., 2016; Takagi et al., 2018)(see Table S2). Clones were selected based on resistance to 100 μg/ml hygromycin B, and Double-AID clones were identified by monitoring the expression of fluorescent markers (e.g., mClover) under a fluorescence microscope. Finally, bi-allelic knock-in of Rad21-mAID-mClover was confirmed by PCR amplification of genomic DNA using KOD FX Neo (KFX-201, TOYOBO, Osaka, Japan) and appropriate primers (Table S3). A parallel attempt to establish Double-AID cell lines enabling double depletion of condensin I and cohesin was unsuccessful, likely due to synthetic lethality caused by leaky depletion of the target subunits. AID cell lines expressing CENP-A-mClover were established using the same protocol. Donor plasmids targeting the CENP-A locus (pMT830 or pMT870) were transfected into H-AID#1, H2-AID#2, Rad21-AID#1, and Double-AID#1 (Table S2). Selection was performed with 100 μg/ml hygromycin B (10687-010, Thermo Fisher Scientific, Waltham, MA, USA) for the first two cell lines, and with 10 mg/ml blasticidin S (A11139-03, Gibco, Thermo Fisher Scientific, Waltham, MA, USA) in the latter two. For these cell lines, we established mono-allelic clones in which one allele was successfully edited as designed. Despite mono-allelic integration, the fluorescence signal from CENP-A-mClover was sufficiently strong for clear visualization of centromeres under the microscope (e.g. Fig. S2, A and D).

### Cell culture

All cells were cultured at 37°C in a humidified atmosphere containing 5% CO₂ in McCoy’s 5A medium (SH30200.01, HyClone, Cytiva, Marlborough, MA, USA) supplemented with 10% heat-inactivated fetal bovine serum (173012, Sigma-Aldrich, St. Louis, MO, USA) and 2 mM L-glutamine (25030-081, Gibco, Thermo Fisher Scientific, Waltham, MA, USA). For fixed-cell immunofluorescence and FISH assays, cells were seeded on poly-L-lysine–coated coverslips (P8920, Sigma-Aldrich, St. Louis, MO, USA). For live-cell imaging, cells were grown on glass-bottom dishes (3910-035, IWAKI, Tokyo, Japan).

To enrich for early G1 cells (Fig. 1 C), cells were first synchronized at the G1/S boundary by treatment with 2.5 mM thymidine (T1895, Sigma-Aldrich, St. Louis, MO, USA) for 16 hr. After release into fresh medium for 5 hr, cells were arrested in G2 phase by treatment with the CDK1 inhibitor RO-3306 (4181/10, Tocris, Minneapolis, MN, USA) at a final concentration of 9 μM for 3 hr (Vassilev et al., 2006). Following washout of RO-3306, cells progressed through mitosis into early G1, during which indole-3-acetic acid (IAA, 19119-61, Nacalai Tesque, Kyoto, Japan) at a final concentration of 0.5 mM or dimethyl sulfoxide (DMSO, D2650, Sigma-Aldrich, St.

Louis, MO, USA) as control (-IAA) was added to deplete the target subunits. Cells were fixed 3 hr after the addition of IAA or DMSO. To enrich for G2-arrested cells (Fig. 4B), cells were treated with 2.5 mM thymidine and released as described above, followed by incubation with the CDK1 inhibitor RO-3306 at a final concentration of 9 μM for 5 hr to induce G2 arrest (Fig. 4 B). IAA or DMSO (no IAA) was added during the final 3 hr of treatment. In experiments shown in Figs. 6 A and S8 A, the duration of RO-3306 and IAA treatments was extended by additional 2 hr.

### Immunofluorescence (IF) assay

Cells cultured on round coverslips (No.1, 12 mm, Matsunami Glass, Osaka, Japan) were fixed with 3.7% paraformaldehyde (PFA, 162-16065, Wako Pure Chemical Industries, Osaka, Japan) or with 3.7% PFA containing 0.5% Triton X-100 (161-11805, Wako Pure Chemical Industries, Osaka, Japan) in PBS (pH 7.5, 1102P05, Cell Science & Technology Institute Inc., Sendai, Japan) for 15 min, followed by permeabilization with 0.5% Triton X-100 in PBS for 5 min at room temperature. After fixation, cells were processed for immunolabeling as described previously (Ono et al., 2013; Ono et al., 2017). Antibodies used in this study are listed in Table S4. A lab-made rabbit polyclonal antibody against human Rad21 (Gandhi et al., 2006) was used for labeling cohesin. A mouse monoclonal antibody, mAb3-19, was used for labeling human CENP-A (Perpelescu et al., 2009). Commercially available antibodies used in this study included mouse monoclonal anti-human Rad21 (53A303; Millipore-Sigma, Burlington, MA, USA), rat monoclonal anti-human CAP-H2 (5F2G4, Active Motif), and rabbit polyclonal anti-human nucleolin (ab22758, Abcam, Cambridge, UK). The following antibodies were used as secondary antibodies: goat anti-rat IgG Alexa Fluor 568 (A11033), goat anti-mouse IgG Alexa Fluor 647 (A21235), goat anti-rabbit IgG Alexa Fluor 488 (A11001), and goat anti-rabbit IgG Alexa Fluor 568 (A11036)(Invitrogen, Thermo Fisher Scientific, Carlsbad, CA, USA). After immunolabeling, DNA were stained with 2 μg/ml DAPI (D1388, Sigma-Aldrich, St. Louis, MO, USA) for 10 min, and the coverslips were mounted with Vectashield H-1000 (Vector Laboratories, Burlingame, CA, USA).

### Fluorescence in situ hybridization (FISH) and IF-FISH assays

Cells cultured on square coverslips (No.1, 18 x 18 mm, Matsunami Glass, Osaka, Japan) were fixed with 2% PFA in 0.3x PBS (pH 7.4) for 10 min. This was followed by extraction with 0.5% Triton X-100 in extraction buffer (20 mM Hepes-KOH [pH 7.9], 50 mM NaCl, 3 mM MgCl2, 300 mM Sucrose) for 10 min, and a second fixation step with 2 % PFA in PBS for 10 min. Then, cells were treated with 0.1 N or 0.2 N HCl for 10 min, and subjected to dehydration through a graded ethanol series. After drying, five μl of probe solution in hybridization buffer (see below) was applied onto the coverslip, which was then mounted on a slide glass (S2111, Matsunami Glass, Osaka, Japan), and sealed with rubber cement (Fixogum, LK-071A, Leica Microsystems, Wetzlar, Germany). The samples were denatured at 80°C for 5 min, followed by hybridization at 37°C for 24 hr in a humidified incubator. After hybridization, cells were washed with 0.4x SSC at 70°C for 2 min, followed by a wash with 2x SSC containing 0.05% Tween-20 (28353-85, Nacalai Tesque, Kyoto, Japan) at room temperature for 5 min.

Subsequently, cells washed once each with 2× SSC, 4× SSC, and PBS for 5 min. DNA were stained with 2 μg/ml DAPI for 10 min and then mounted with Vectashield H-1000. For IF-FISH experiments, cells were first fixed and processed for immunofluorescence using an anti-nucleolin antibody (Table S4). Subsequently, cells were fixed again with 2% PFA in PBS for 10 min. Hybridization and all subsequent procedures were performed as described for the FISH experiments.

All conventional FISH probes (Table S5) were dissolved in hybridization buffer and provided as ready-to-use solutions (MetaSystem probes, Altlussheim, Germany). These probes were used in combination, and their mixing ratios are listed in Table S7. For arm conformation analysis, oligo probes were also dissolved in the same hybridization buffer (Table S7). These oligo probes consisted of pools of 43-47 nt long single-stranded DNAs covering a ∼300 kb of target regions (myTags, Primetech/Arbor Biosciences, MI, USA)(Table S6). They were designed at a density of ∼10 probes per kilobase (kb) and labeled with different fluorescent dyes.

### Microscopy

Fluorescence images were captured using a DeltaVision Elite system (Cytiva, Marlborough, MA, USA, formerly Applied Precision) equipped with an inverted microscope (IX71, Olympus, Tokyo, Japan). UPlanSApo 100×/1.40 NA and UPlanXApo 100×/1.45 NA oil-immersion objective lenses (Olympus, Tokyo, Japan) were used for immunofluorescence and FISH assays using laser immersion oil (refractive index 1.506, Laser Liquid, Moritex Co., Ltd., Yokohama, Japan).

UPlanSApo 60×/1.30 NA silicone immersion oil objective lenses were used for live imaging using silicon immersion oil (SIL300CS-30CC, refractive index 1.406, Olympus, Tokyo, Japan). Image stacks of entire cells were acquired with a Z-step size of 0.3 μm for immunofluorescence, 0.5 μm for FISH and IF-FISH, and 2 μm for live imaging. Z-stack images were processed by constrained iterative deconvolution (10 iterations) as described previously (Chen et al., 1996). For immunofluorescence and FISH using WCP probes, 7 to 30 sections were projected as average-intensity projections. For FISH using site-specific probes and live imaging, maximum-intensity projections were used. Grayscale images were pseudocolored and merged using Photoshop (Adobe, San Jose, CA).

### Quantification of immunofluorescent intensity and centromere dispersion

Quantitative analysis of immunofluorescent signals was performed using Image J (https:// imagej.net/ij/index.html). For nuclear intensity measurements of CAP-H2 and Rad21, nuclei were segmented by thresholding the DAPI signal, and the mean gray values of CAP-H and Rad21 signals within each nuclear region were measured. After background subtraction, total fluorescence intensities per nucleus were plotted using GraphPad Prism8 (GrapPad Software, San Diego, CA, USA)(Figs. 1, D-F, S2 F, S4, and S7). Statistical significance was assessed using the Mann-Whitney U test. To assess centromere dispersion, we calculated a “dispersion index”, defined as the area of the convex hull (the smallest convex polygon enclosing all CENP-A signals) divided by the DAPI-positive nuclear area. Using uniformly binarized images of CENP-A, the convex hull of the signals was manually delineated. Dispersion indices in fixed cells and live cells were plotted using GraphPad Prism8 Software (Figs. 1 G and S2 B) and Microsoft Excel (Fig. S2 E), respectively. Statistical significance (P value) of dot plots was assessed using the Kruskal-Wallis test.

### Quantification of area, distance, morphology and contact ratio in FISH/IF-FISH

Analysis of nuclear and CT areas was performed as described above. DAPI-and FISH-positive signal areas were segmented by thresholding and plotted (Figs. 3 F, S2, B and G, and S3, D and H). Chromosome segmentation in FISH images was performed using individual thresholding to each image to account for cell-to-cell variations in signal intensity. To analyze the morphology of nuclei, CTs, and nucleoli, we used “circularity“, a morphological index that indicates how closely the shape of an object approximates a perfect circle (with a value of 1 indicating a perfect circle), calculated from the segmented regions. Circularity values were plotted using GraphPad Prism8 Software (Figs. 3 G, and 6, G and H). Statistical significance (P value) for dot plots and bar graphs was assessed using the Kruskal-Wallis test, and the Chi-square test, respectively.

For the analysis of distances between paired signals (Figs. 2 E, 4 F, and S5, A and B), the relevant FISH signals were segmented by thresholding, and the coordinates of each signal centroid were determined. Distances between paired signals was calculated using these coordinates and standard trigonometric functions, and plotted accordingly (Figs. 2 E, 4 F, and S5, A and B). In principle, two signals are expected at each locus in G2-phase nuclei, as sister chromatids have already replicated. However, sister signals were often indistinguishable in our observations, likely due to close proximity or signal overlap. In such cases, the same distance value was assigned to both chromatids. Statistical significance (P value) for dot plots and bar graphs was assessed using the Kruskal-Wallis test, and the Chi-square test, respectively.

To evaluate CT-nucleolus contact, we quantified the “contact ratio”, defined as the length of the CT perimeter in contact with the nucleolus divided by the total CT perimeter. Chromosome 18 territories and nucleoli were segmented by thresholding, and CT perimeters were measured using ImageJ. The length of the interface where chromosome 18 was in contact with the nucleolus was manually measured using the line tool in ImageJ. The resulting contact ratios were plotted using GraphPad Prism8 Software (Fig. 6 F). Statistical significance (P value) for dot plots and bar graphs was assessed using the Kruskal-Wallis test, and the Chi-square test, respectively.

### Quantification of angular distribution

To assess chromosome arm conformation, we defined the “P angle” as the angle formed between the lines connecting point P to D3 and point P to C, which reflect the degree of arm bending (Fig. 5 B). Three FISH signal regions were segmented by thresholding, and the centroid coordinates of each region were determined using ImageJ. Given the coordinates of the three points forming a triangle, the internal angles at each vertex were calculated using the law of cosines. Among these, the P angles were extracted and plotted (Figs. 5, C-E, and S6, B-C and E-H). To test for angular bias in datasets where angles ±θ are considered equivalent (symmetric around 0°), each observed angle θ_j was duplicated into two values, +θ_j and −θ_j. To assess differences in angular distributions between two groups, we used the Mardia-Watson-Wheeler test (Mardia, 1967; Kondo and Kimura, 2019). The Rayleigh test for uniformity (Fisher, 1995) was then applied to the expanded dataset using the resultant vector length R. The test statistic was computed as Z = N R^2^, where N represents the number of original (independent) angle measurements, not the total number after duplication. A significant Z-value indicates deviation from a uniform angular distribution. (Figs. 5 F and S6 I).

## Acknowledgments

We thank M. Kanemaki and T. Natsume (Division of Molecular Cell Engineering, NIG, Japan) for providing the Rad21-AID cell line and plasmid constructs. We are also grateful to the late K. Yoda (Nagoya Univ.) for the CENP-A antibody and to K. Ohtawa (RIKEN) for technical assistance with FACS analysis. We thank the members of the Hirano laboratory for their critical reading of the manuscript. This work was supported by the Japan Society for the Promotion of Science (JSPS) KAKENHI (Grant Number 21K06027 [to T.O.]; 18H05276 and 20H05938 [to T.H.]).

## Author contributions

Author contributions: T. Ono: Conceptualization, Data curation, Formal analysis, Funding acquisition, Investigation, Methodology, Project administration, Validation, Visualization, Writing - original draft, Writing - review & editing. M. Takagi: Formal analysis, Methodology, Resources. H. Tanabe: Methodology. T. Fujita: Formal analysis, Methodology, N. Saitoh: Formal analysis, Methodology. A. Kimura: Formal analysis, Methodology. T. Hirano: Conceptualization, Funding acquisition, Project administration, Supervision, Writing - original draft, Writing - review & editing.

## Supplemental figure legends

**Figure S1.**
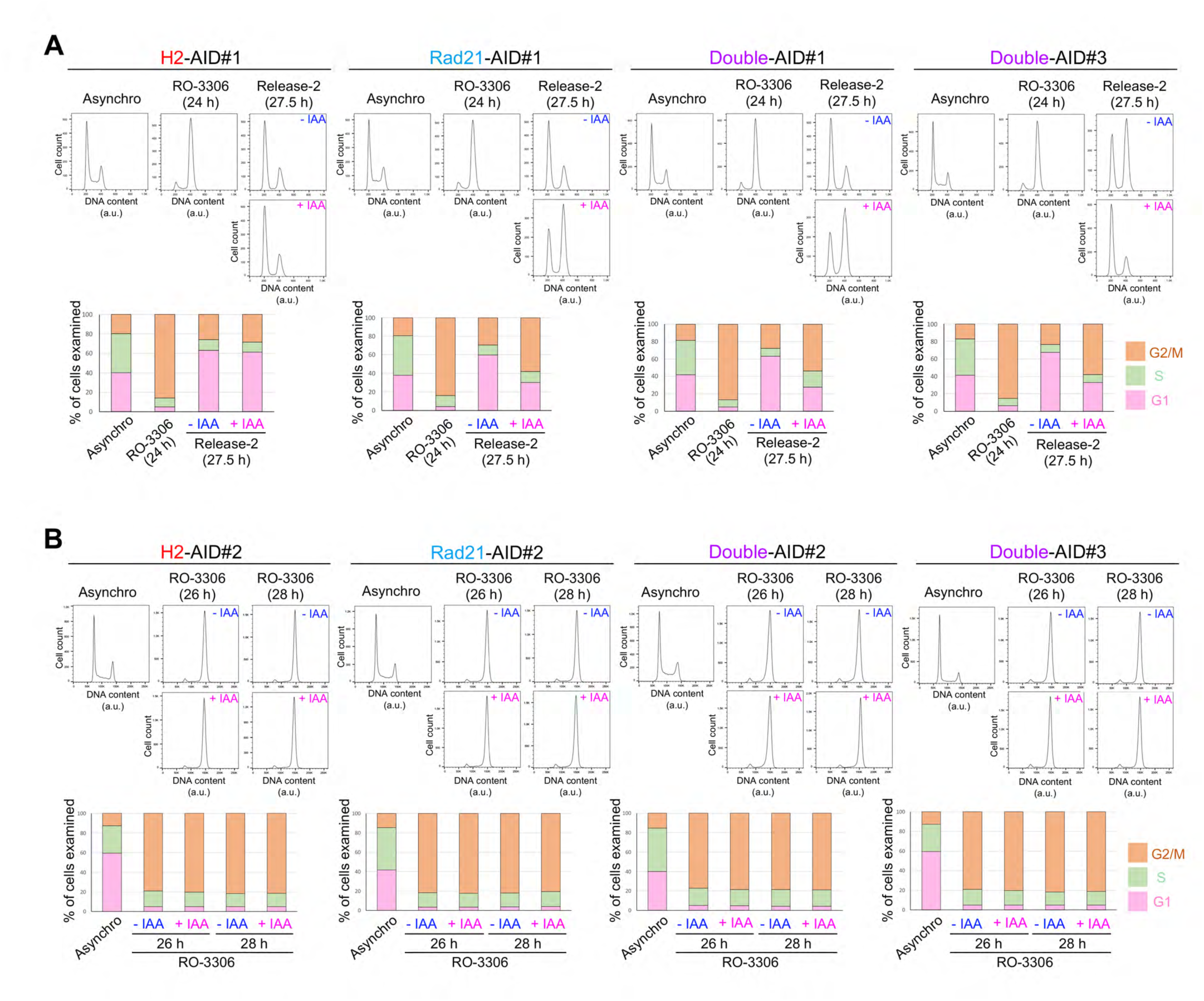
Assessment of cell cycle synchronization by FACS analyses. (Related to. **Figures 1 C, 4 A, and 6 A)** For FACS analysis, cells were washed twice with PBS and fixed with cold 70% ethanol. After incubation with 10 μg/ml RNase and 50 μg/ml propidium iodide at 37°C for 30 min, cells were analyzed using a flow cytometer. **(A)** Cells undergoing the M-to-G1 transition were prepared according to the protocol shown in Fig. 1 C, and analyzed using a flow cytometer (LSR, BD). Fluorescence signals were collected at a wavelength of 575 nm, and the data were analyzed using FlowJo software (Tree Star). In this protocol, ∼60% of control and H2-depleted cells, and ∼30% of Rad21-depleted and co-depleted cells, were successfully synchronized in G1 phase. The apparently lower synchronization efficiency in the latter two groups is attributable to the well-documented mitotic delay caused by cohesin depletion (Hauf et al., 2005; Haarhuis et al., 2013; Perea-Resa et al., 2020). From these synchronized populations, early G1 cells were selected based on their characteristic morphologies (see the legend of Fig. 1 C). **(B)** G2-arrested cells were prepared according to the protocol shown in Figs. 4 A and 6 A, and analyzed using a flow cytometer (A3, BD). Fluorescence signals were collected at a wavelength of 610/20 nm, and the data were analyzed using BP FlowJo software (BD). In this protocol, ∼80% of cells were successfully synchronized in G2 phase across all cell lines. Extending the duration of IAA treatment from 3 hr to 5 hr did not affect the efficiency of G2 synchronization (Figs. S4 and S7).

**Figure S2.**
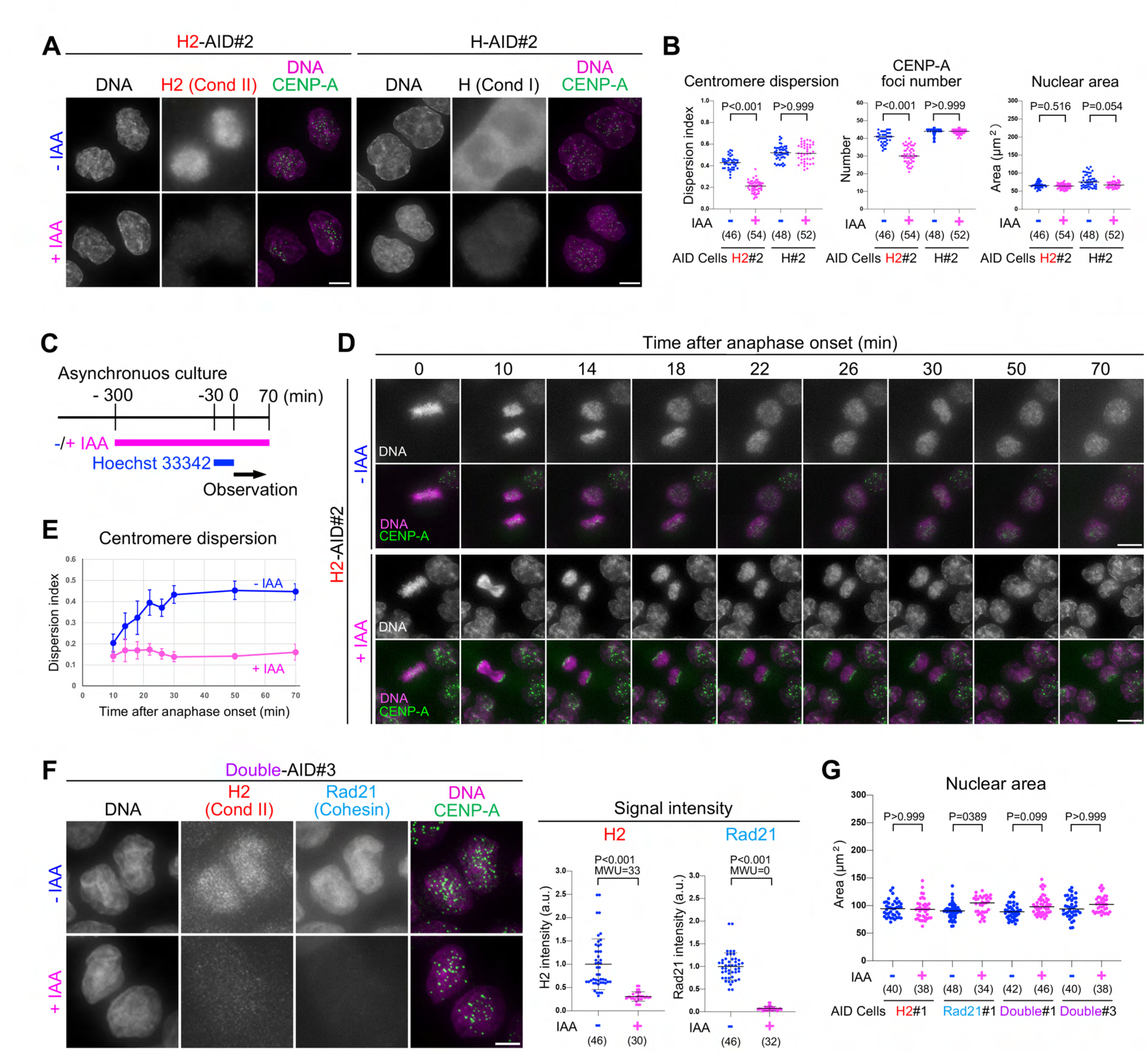
Additional analyses of M-to-G1 transition phenotypes. (Related to Figure 1) **(A)** Condensin I depletion does not suppress post-mitotic centromere dispersion. For this analysis, H-AID#2 and H2-AID#2 cells, both expressing CENP-A-mClover, were used. Cells cultured asynchronously on coverslips were treated with (+) or without (-) IAA for 5 hr, and then fixed with 3.7% PFA in PBS, followed by extraction with 0.5% Triton X-100 in PBS. DNA was counterstained with DAPI. Bar, 5 μm. **(B)** Dot plots show the dispersion index of nuclear CENP-A foci, the number of CENP-A foci, and the nuclear area. The definition of the dispersion index is shown in Fig. 1 G. The numbers in parentheses at the bottom of the plot indicate the number of cells examined. Bars represent median values. The same set of images was used for the three quantitative analyses under each condition. **(C)** Schematic diagram of the cell preparation protocol for live-cell imaging. **(D)** Asynchronously cultured H2-AID#2 cells were treated with (+) or without (-) IAA for 5 hr, and then processed for live imaging. The first frame at anaphase onset was defined as time point 0 (min). Bar, 10 μm. **(E)** Plots showing time-course changes in the dispersion index as defined in Fig. 1 G. Bars represent the mean ± sd values (n = 6). **(F)** Centromere dispersion (left) and quantification of nuclear signal intensities of CAP-H2 and Rad21(right) in an alternative cell line, Double-AID#3. As shown in Fig. 1, E and F, fixed cells were immunolabeled with an anti-CAP-H2 antibody. DNA was counterstained with DAPI. Bar, 5 μm. Dot plots show the nuclear signal intensities of CAP-H2 (IF) and Rad21-mClover. The numbers in parentheses at the bottom of the panel indicate the number of cells analyzed. P, p-value. MWU, the Mann-Whitney U test statistic. Bars represent the mean ± sd values. a.u., arbitrary unit. **(G)** Dot plots show the nuclear area of G1 cells analyzed for CENP-A in Figs. 1, D–F, and S2 F. Bars represent median values. The same set of FISH images analyzed in Fig. 1, G and H was used for the analysis.

**Figure S3.**
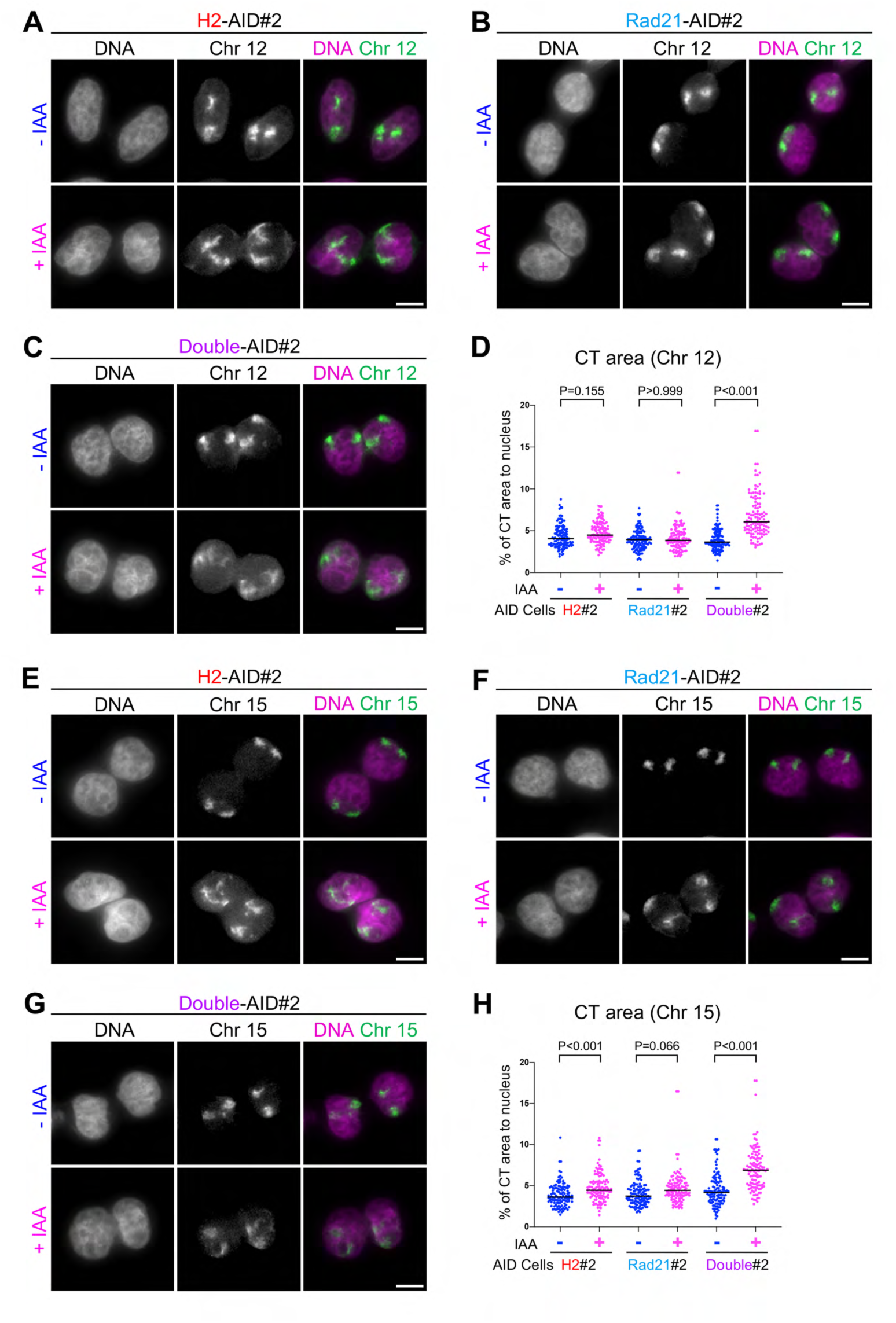
Chromosomes 12 and 15 also exhibit severe defects in CT morphology in G1 cells co-depleted of condensin II and cohesin. (Related to. **Figures 2 and 3) (A-C)** H2-AID#2 (A), Rad21-AID#2 (B), and Double-AID#2 (C) cells were treated as described in Fig. 3 B, fixed, and processed for FISH using a WCP probe for chromosome 12 (Table S5). DNA was counterstained with DAPI. Bar, 5 μm. **(D)** Dot plots show the area of FISH-labeled territories for chromosomes 12. Bars represent median values. **(E-G)** H2-AID#2 (E), Rad21-AID#2 (F), and Double-AID#2 (G) cells were treated as described in (A-C), fixed, and processed for FISH using a WCP probe for chromosome 15 (Table S5). DNA was counterstained with DAPI. Bar, 5 μm. **(H)** Dot plots show the area of FISH-labeled territories for chromosomes 15. Bars represent median values. Panels (D) and (H) were analyzed with 120 chromosomes (from 60 cells) in each condition.

**Figure S4.**
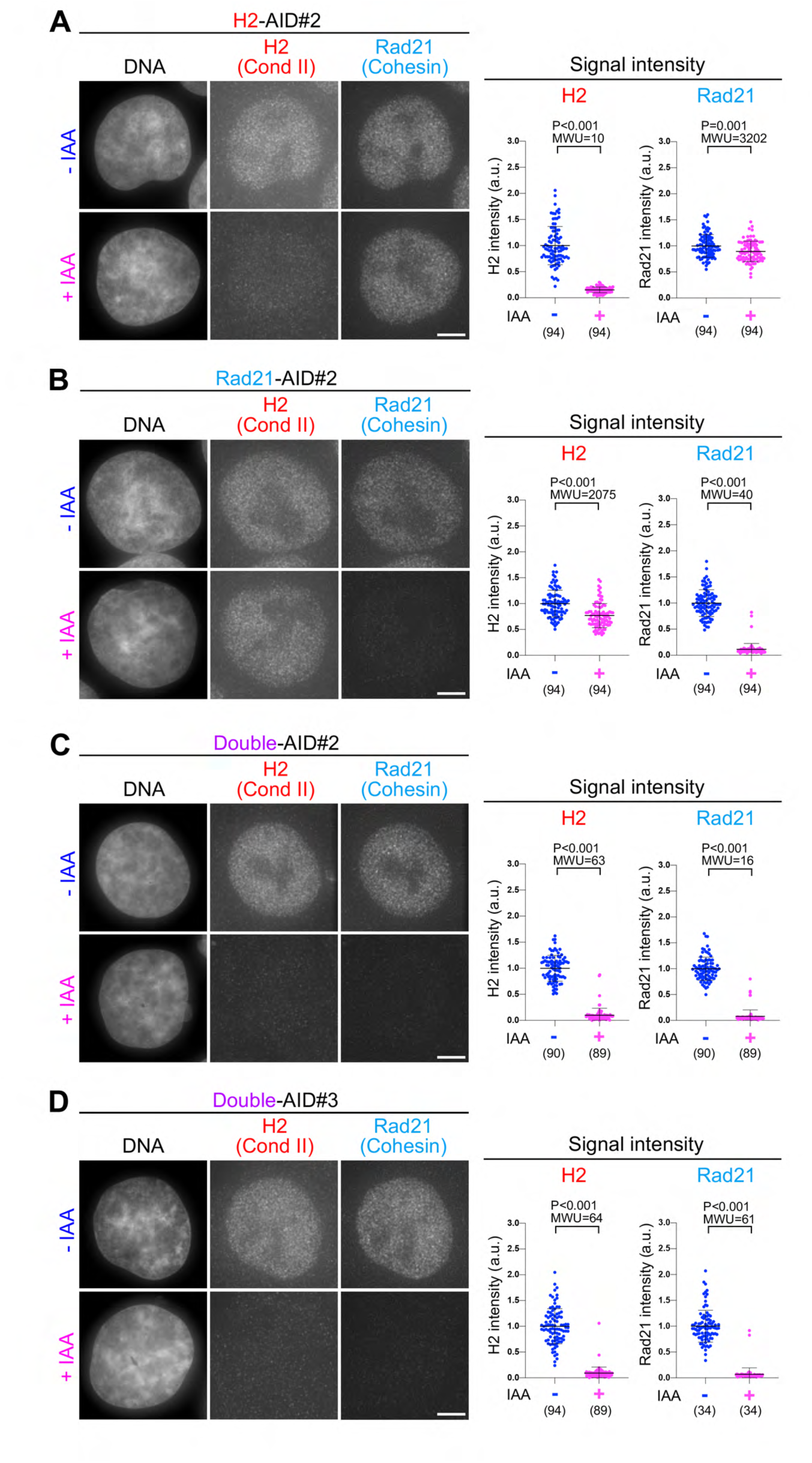
Depletion of condensin II and/or cohesin in G2-arrested cells. (Related to Figures 4 and 5) (**A-D**) H2-AID#2 (A), Rad21-AID#2 (B), Double-AID#2 (C) and Double-AID#3 (D) cells were treated according to the protocol described in Fig. 4 B. Cells were fixed with 3.7% PFA in PBS containing 0.1% Triton X-100, extracted with 0.5% Triton X-100 in PBS, and then immunolabeled with antibodies against CAP-H2 and Rad21. DNA was counterstained with DAPI. Scale bar, 5 μm. Dot plots show the nuclear signal intensities of CAP-H2 and Rad21. The numbers in parentheses at the bottom of the panel indicate the number of cells analyzed. P, p-value. MWU, the Mann-Whitney U test statistic. Bars represent the mean ± sd values. a.u., arbitrary unit.

**Figure S5.**
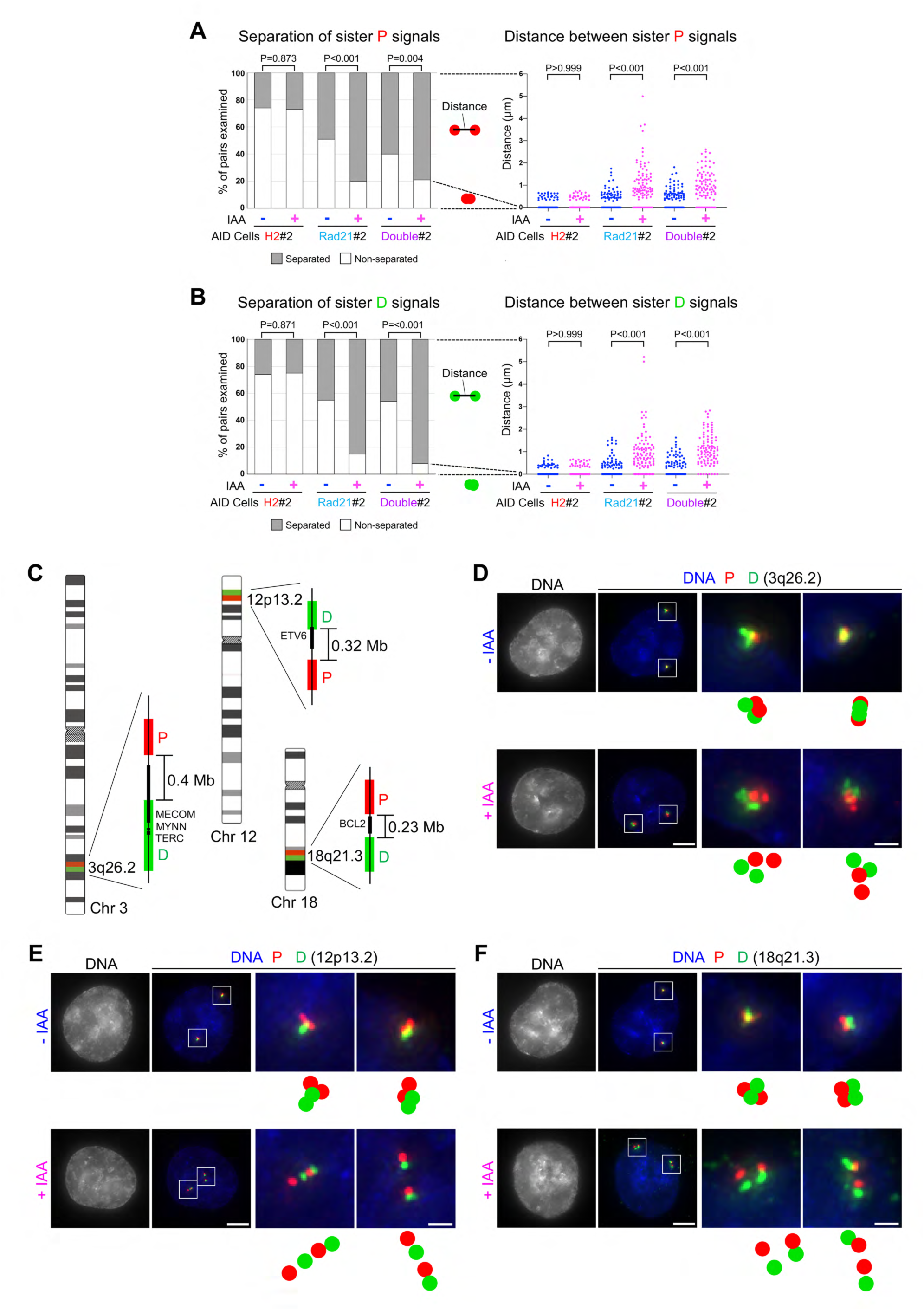
Additional analyses of G2-arrested phenotypes. (Related to. **Figure 4)** (**A and B**) The same set of FISH images analyzed in Fig. 4, C-E was used for the analysis of sister chromatid resolution presented here. Bar graphs (left) show the proportions of signal pairs with separated sister P signals (A) and sister D signals (B). Gray bars indicate distinguishable sister signals, whereas white bars indicate indistinguishable ones. Dot plots (right) show the distribution of distances between distinguishable sister P signals (A) and distinguishable sister D signals (B). Bars represent median values. **(C)** Additional FISH probes used to analyze local chromatin folding. We employed three pairs of site-specific FISH probes targeting at 3q26.2, positioned 0.4 Mb apart, 12q13.2, positioned 0.32 Mb apart, and 18q21.3 positioned 0.26 Mb apart (Table S5). **(D-F)** Rad21-AID#2 cells were treated according to the protocol described in Fig. 4 B, fixed, and processed for FISH using probes targeting at 3q26.2 (D), 12q13.2 (E), and 18q21.3 (F). DNA was counterstained with DAPI. Bar in the left panels, 5 μm. The regions enclosed by the white rectangles are magnified and shown on the right. Pairs of P (proximal; red) and D (distal; green) judged to be on the same chromosome are connected by black lines in the cartoons shown at the bottom. Among the four loci examined in this study including 11q13.3 (Fig. 4 A), changes observed upon cohesin depletion appeared to be locus-dependent. Notably, the locus on 11q showed the most prominent change, coincided with the loss of a TAD structure as previously reported (Rao et al., 2017). The corresponding Hi-C map can be viewed at: https://3dgenome.fsm.northwestern.edu/index.html. Therefore, we used the 11q locus for comparison with condensin II depletion and double depletion.

**Figure S6.**
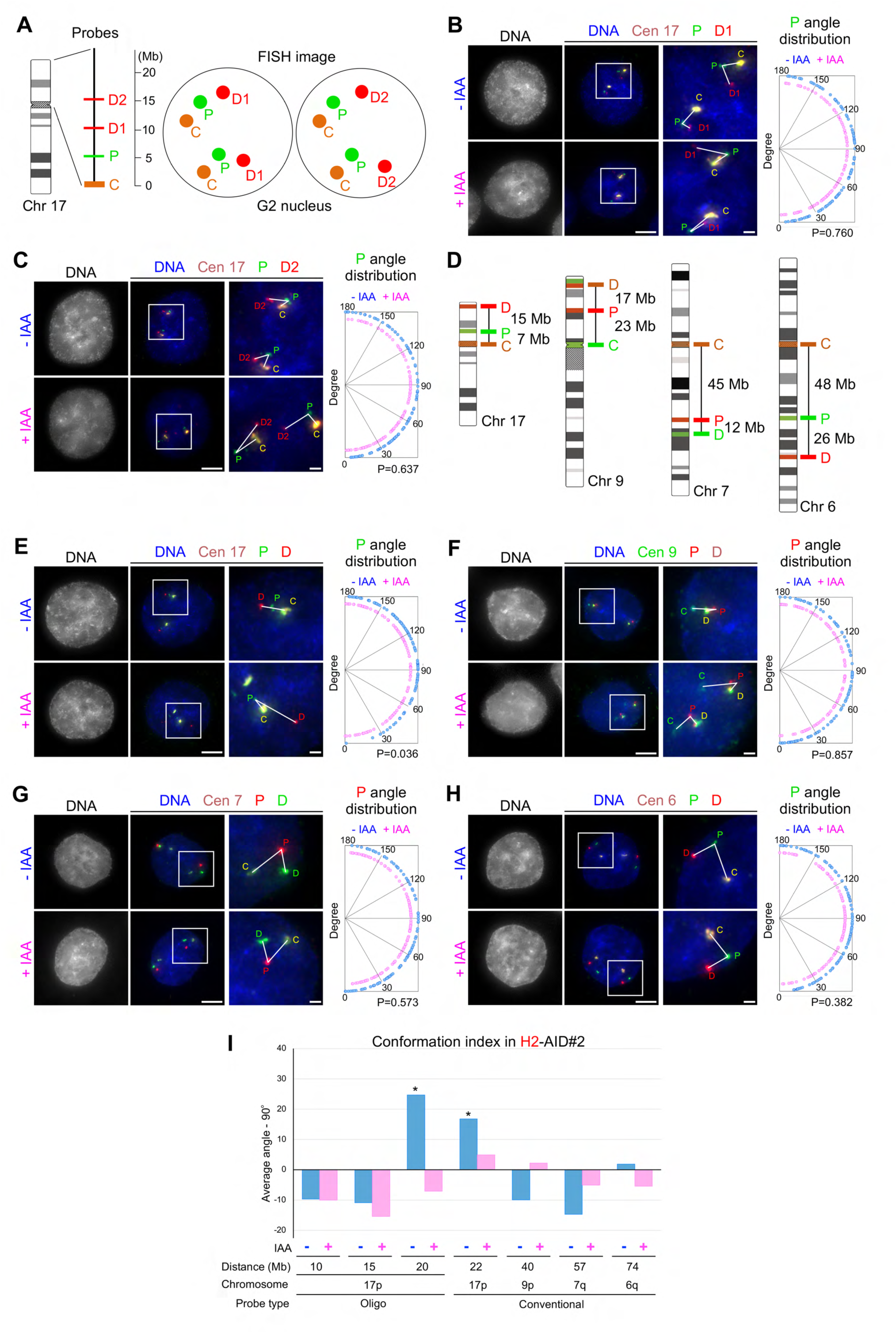
Condensin II has its most prominent effect on chromosome arm conformation at the ∼20 Mb genomic scale. (Related to Figure 5) **(A)** FISH probes used and a schematic of FISH signals expected. A set of three site-specific probes was used: two custom oligo-probes targeting specific loci (P and D1 or D2) on the short arm of chromosome 17, and a mixture of centromere-specific probes for chromosome 17. Oligo probes P, D1, and D2 locates at 5 Mb, 10 Mb, and 15 Mb away from the centromere, respectively (Tables S6 and S7). Consequently, two groups of FISH signals including centromeres (yellow), P sites (green), and D1 sites (red), or centromeres (yellow), P sites (green), and D2 sites (red) are delineated within a nucleus, as depicted in the right panel. **(B and C)** Left: H2-AID#2 cells were treated according to the protocol described in Fig. 4 B, fixed, and processed for FISH using sets of three site-specific probes described in (A). Bar in left panels, 5 μm. The regions enclosed by the white rectangles are magnified and shown in the third columns. Sets of C (yellow), P (green) and D1 (red) judged to be on the same chromosome are connected by white lines. Bar in right panel, 1 μm. Right: P angle distributions in the cells treated with (+) or without (-) IAA are shown as circular scatter plots. Statistical significance was assessed using the Mardia Watson Wheeler test, as described in Fig. 5, C-E. In this experiment, 100 signal sets (i.e. chromosomes) from 50 cells were analyzed under each condition. **(D)** Additional sets of FISH probes used. Each set included a mixture of centromere-specific probes (yellow) and two site-specific probes (green and red)(Tables S5 and S7). Genetic distances between each probes are shown within the cartoon. **(E-H)** Left: H2-AID#2 cells were treated according to the protocol described in (B and C), fixed, and processed for FISH using sets of site-specific probes (D). Bar in left panels, 5 μm. The regions enclosed by the white rectangles are magnified and shown in the third columns. Sets of C (yellow), P (green), and D (red) judged to be on the same chromosome are connected by white lines. Bar in right panel, 1 μm. Right: P angle distribution in cells treated with (+) or without (-) IAA are shown as circular scatter plots. Statistical significance was assessed using the Mardia Watson Wheeler test. In this experiment, 100 signal sets (i.e. chromosomes) from 50 cells were analyzed under each condition. **(I)** Quantification of the P angle in H2-AID#2 cells. The bar graphs show “conformation index”, defined as the average angle P minus 90° (Fig. 5 F). To facilitate comparison with the 20-Mb scale analysis using oligo probes, data from the H2-AID#2 cell line shown in Fig. 5 F were incorporated into this graph. Deviations from randomness were assessed using the Rayleigh test. Asterisks indicate statistically significant deviations from a random distribution (P< 0.05).

**Figure S7.**
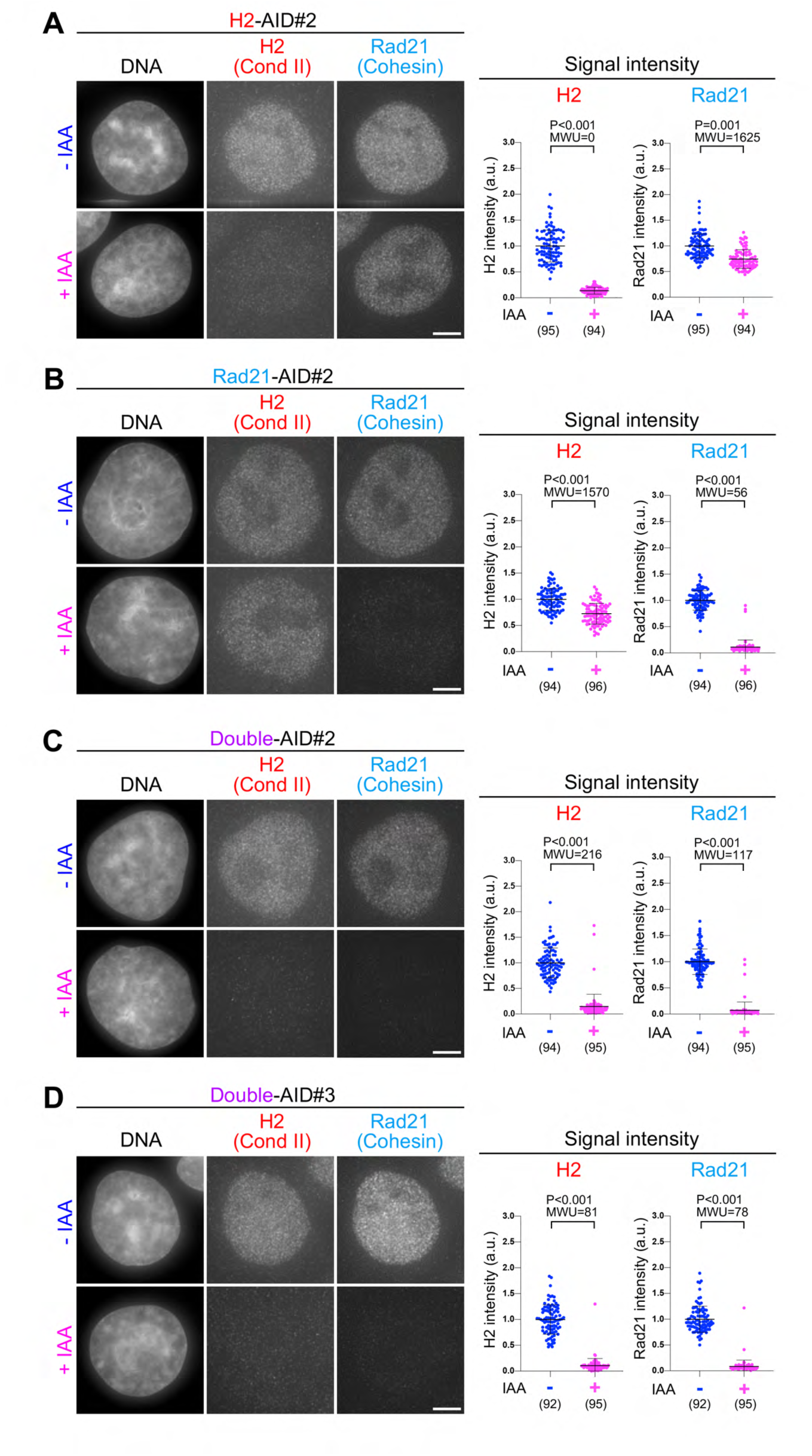
Prolonged incubation does not alter the signal intensities of the target subunits in G2-arrested cells. (Related to Figures 6 and S4) (**A-D**) H2-AID#2 (A), Rad21-AID#2 (B), Double-AID#2 (C) and Double-AID#3 (D) cells were treated according to the protocol described in Fig. 6 A. Cells were fixed, extracted, and then immunolabeled with antibodies against CAP-H2 and Rad21 as described in Fig. S4. DNA was counterstained with DAPI. Scale bar, 5 μm. Dot plots show the nuclear signal intensities of CAP-H2 and Rad21. The numbers in parentheses at the bottom of the panel indicate the number of cells analyzed. P, p-value. MWU, the Mann-Whitney U test statistic. Bars represent the mean ± sd values. a.u., arbitrary unit.

**Figure S8.**
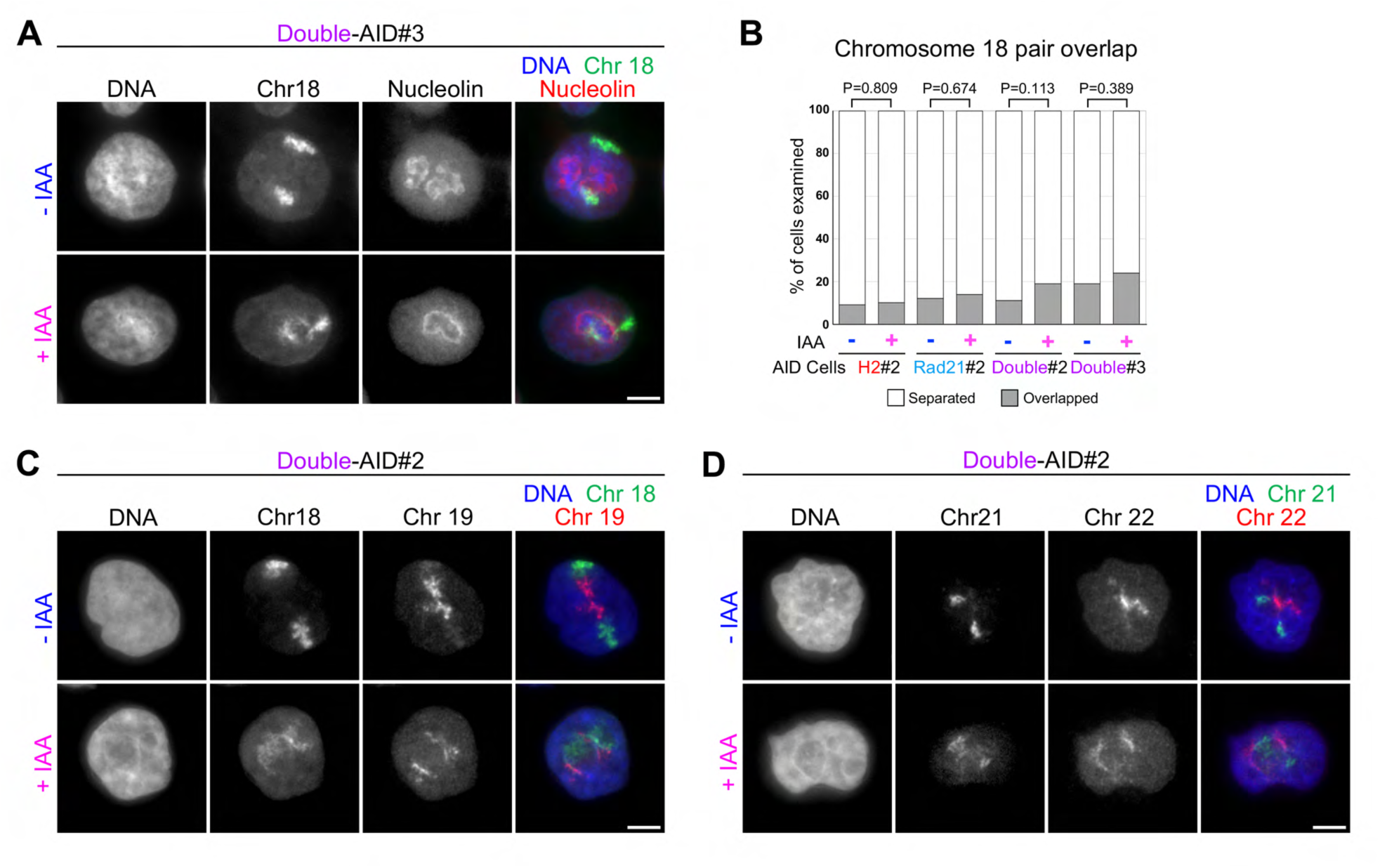
Additional analyses of CT morphology observed in G2-arrested cells co-depleted of condensin II and cohesin. (Related to Figure 6) **(A)** Double-AID#2 cells were treated according to the protocol described in Fig. 6 A. Cells were immunolabeled with an anti-nucleolin antibody, and then processed for FISH using sets of three WCP probes for chromosomes 18 as described in Fig. 6, C-E. Bar, 5 μm. **(B)** Quantification of the overlap between chromosome 18 homologs in four cell lines examined. The same images were used in Fig. 6, C-E and (A), with 100 pairs of chromosome 18 from 100 cells under each condition. **(C and D)** Double-AID#2 cells were treated according to the protocol described in Fig. 2, B-D. After fixation and extraction, cells were processed for FISH using WCP probes for chromosomes 18 and 19 (C), and for chromosomes 21 and 22 (D). Chromosome 19 was deformed and exhibited abnormal perinucleolar distribution (C). Chromosomes 21 and 22 also exhibited deformations similar to those observed for chromosomes 18 and 19 (D). Bar, 5 μm.

---

**Table S1.** AID cell lines used in this study.

| Cell line | Abbreviation | Reference |
| --- | --- | --- |
| CAP-H-mACherry | H-AID#1 | Takagi et al. (2017) |
| CAP-H2-mACherry | H2-AID#1 | Takagi et al. (2017) |
| RAD21-mAID-mClover | Rad21-AID#1 | Natsume et al. (2016) |
| CEP-A-mClover/CAP-H-mACherry | H-AID#2 | this study |
| CEP-A-mClover/CAP-H2-mACherry | H2-AID#2 | this study |
| CEP-A-mClover/RAD21-mAID-mClover | Rad21-AID#2 | this study |
| CAP-H2-mACherry/RAD21-mAID-mClover (13-28) | Double-AID#1 | this study |
| CEP-A-mClover/CAP-H2-mACherry/RAD21-mAID-mClover | Double-AID#2 | this study |
| CAP-H2-mACherry/RAD21-mAID-mClover (13-48) | Double-AID#3 | this study |

**Table S2.**
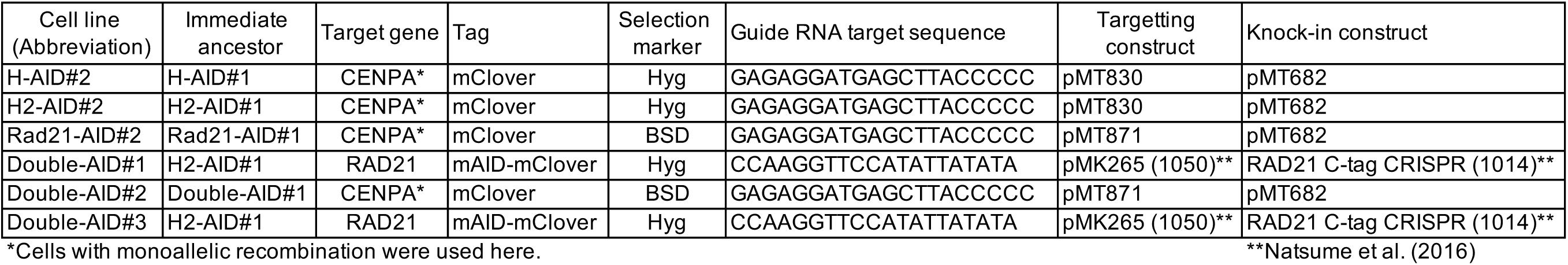
AID cell lines established in this study.

**Table S3.**
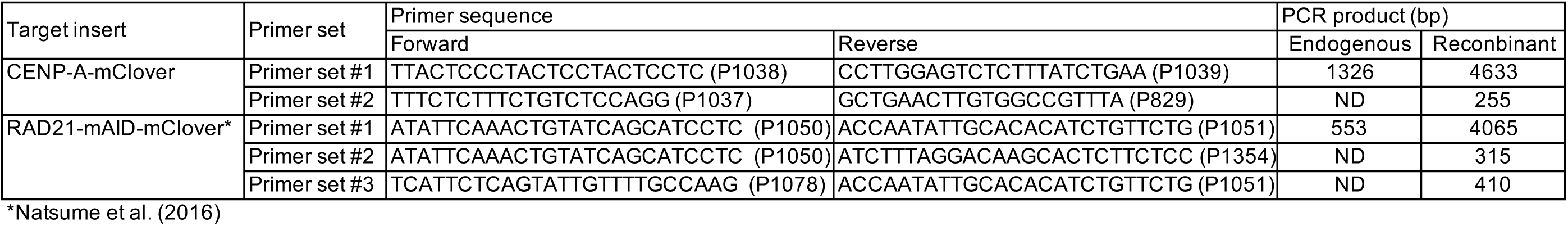
Primers used for the genomic PCR.

**Table S4.** Antibodies used int this study.

| Antigen | Species | Manufacturer or Provider | Clone name or Catalog # | IF final conc. | Figure | Reference |
| --- | --- | --- | --- | --- | --- | --- |
| CAP-H2 | Rat | Active Motif | 5F2G4/61682 | 2µg/mL | Fig. 1, B and D-F; Fig. S2 F; Fig. S4, A-D; Fig. S7, A-D |  |
| Rad21 | Rabbit | Hirano Lab | AfR229-2L | 2µg/mL | Fig. 1 D | Gandhi, et al. (2006) |
| Rad21 | mouse | Millipore | 53A303/m05-908 | 1µg/mL | Fig. 1 B; Fig. S4, A-D; Fig. S7, A-D |  |
| CENP-A | mouse | Provided by K. Yoda | mAb 3-19 | 1:4 | Fig. 1, D-F; Fig. S2 F | Perpelescu et al. (2009) |
| Nucleolin | Rabbit | Abcam | ab22758/RB568 | 1µg/mL | Fig. 6, C-E; Fig. S8 A |  |

All secondary antibodies conjugated to Alexa Fluor 488, 568, or 647 (Invitrogen, Thermo Fisher Scientific) were used at a 1:500 dilution (See Materials and Methods).

**Table S5.** FISH probes used int this study.

| Conventional probe (Manufacturer: MetaSystem probes) |  |  |  |
| --- | --- | --- | --- |
| Name | Target | Description or Catalog # | Figure |
| XCP18 green | Whole chromosome 18 | D-0318-050-FI | Fig. 3, B-D; Fig. 6, C-E; Fig.S8, A and C |
| XCP19 orange | Whole chromosome 19 | D-0319-050-OR | Fig. 3, B-D; Fig.S8 C |
| XL MDM2 | Cen12 (green) and 12q15 (orange) | D-5047-100-OG | Fig. 2, B-D |
| XL CCND1 | Two loci (green and orange) in 11q13.3 (0.56 Mb apart) | D-5071-100-OG | Fig. 4, C-E |
| XCE17 green | Cen 17 | D-0817-050-FI | Fig. 5, C-E; Fig. S6, B, C, and E |
| XCE17 orange | Cen 17 | D-0817-050-OR | Fig. 5, C-E; Fig. S6, B, C, and E |
| XCP12 green | Whole chromosome 12 | D-0312-050-FI | Fig. S3, A-C |
| XCP15 green | Whole chromosome 15 | D-0315-050-FI | Fig. S3, E-G |
| XL MECOM 3q26 | Two regions (green and orange) in 3q26 (0.4 Mb apart) | D-5059-100-OG | Fig. S5 D |
| XL ETV6 | Two regions (green and orange) in 12q13.2 (0.32 Mb apart) | D-5073-100-OG | Fig. S5 E |
| XL BCL2 BA | Two regions (green and orange) in 18q21.3 (0.26 Mb apart) | D-6018-100-OG | Fig. S5 F |
| XL Smith-Magenis/Miller-Dieker | 17p11.2 (green), 17p13 (orange) | D-5422-050-OG | Fig. S6 E |
| XCE 9 green | Cen 9 | D-0809-050-FI | Fig. S6 F |
| XL CDKN2A | Cen 9 (green), 9p21 (orange) | D-5053-100-OG | Fig. S6 F |
| XL JAK2 BA | 9p24 (green) and 9p24 orange) | D-5098-100-OG | Fig. S6 F |
| XCE 7 green | Cen 7 | D-0807-050-FI | Fig. S6 G |
| XCE 7 orange | Cen 7 | D-0807-050-OR | Fig. S6 G |
| XL del(7)(q22q31) | 7q22 (orange) and 7q31.2 (green) | D-5068-100-TC | Fig. S6 G |
| XCE 6 green | Cen 6 | D-0806-050-FI | Fig. S6 H |
| XCE 6 orange | Cen 6 | D-0806-050-OR | Fig. S6 H |
| XL 6q21/6q23 | 6q21 (green), 6q23 (orange) | D-5039-100-OG | Fig. S6 H |
| XCP21 green | Whole chromosome 21 | D-0321-050-FI | Fig. S8 D |
| XCP22 orange | Whole chromosome 22 | D-0322-050-OR | Fig. S8 D |

**Table S6.** Oligo probes used int this study.

| myTag Oligo probe (Manufacturer: Primetech/Arbor Bioscience) |  |  |  |  |  |
| --- | --- | --- | --- | --- | --- |
| Name | Region | Count | Ave density* | Label | Figure |
| Chr 17 P | chr17:19800000-20100000 | 3196 | 10.67 | Alexa-488 | Fig. 5, C-E; Fig. S6, B and C |
| Chr 17 D1 | chr17:14800000-15100000 | 4242 | 14.14 | ATTO-565 | Fig. S6 B |
| Chr 17 D2 | chr17:9800000-10100000 | 2785 | 9.28 | ATTO-565 | Fig. S6 C |
| Chr 17 D3 | chr17:5171000-5471000 | 3058 | 10.26 | ATTO-565 | Fig. 5, C-E |

\*Probes/kb

**Table S7.** Probe composition for multicolor FISH.

| Target | Probe | Probe type | Ratio or concentration | Figure |
| --- | --- | --- | --- | --- |
| Chr 18 and Chr 19 | XCP18 green | Conventional | Volume ratio of 1:1 | Fig. 3, B-D; Fig. S8 C |
|  | XCP19 orange | Conventional |  |  |
| Cen 17, P, and D1-D3 | XCE17 green | Conventional | Volume ratio of 1:1 | Fig. 5, C-E; Fig. S6, B and C |
|  | XCE17 orange | Conventional |  |  |
|  | P | Oligo | 0.5 pm/μL |  |
|  | D1, D2, or D3 | Oligo | 0.5 pm/μL |  |
| Cen 17, P, and D | XCE17 green | Conventional | Volume ratio of 3:3:5 | Fig. S6 E |
|  | XCE17 orange | Conventional |  |  |
|  | XL Smith-Magenis/Miller-Dieker | Conventional |  |  |
| Cen 9, P, and D | XCE 9 green | Conventional | Volume ratio of 3:4:4 | Fig. S6 F |
|  | XL CDKN2A | Conventional |  |  |
|  | XL JAK2 BA | Conventional |  |  |
| Cen 7, P, and D | XCE 7 green | Conventional | Volume ratio of 3:3:5 | Fig. S6 G |
|  | XCE 7 orange | Conventional |  |  |
|  | XL del(7)(q22q31) | Conventional |  |  |
| Cen 6, P, and D | XCE 6 green | Conventional | Volume ratio of 3:3:5 | Fig. S6 H |
|  | XCE 6 orange | Conventional |  |  |
|  | XL 6q21/6q23 | Conventional |  |  |
| Chr 21 and Chr 22 | XCP21 green | Conventional | Volume ratio of 1:1 | Fig. S8 D |
|  | XCP22 orange | Conventional |  |  |

## References

Abramo, K., A.L. Valton, S.V. Venev, H. Ozadam, A.N. Fox, and J. Dekker. 2019. A chromosome folding intermediate at the condensin-to-cohesin transition during telophase. Nat. Cell Biol. 21:1393–1402. doi: [10.1038/s41556-019-0406-2].

Batty, P., and D.W. Gerlich. 2019. Mitotic Chromosome Mechanics: How Cells Segregate Their Genome. Trends Cell Biol. 29:717–726. doi: [10.1016/j.tcb.2019.05.007].

Bauer, C.R., T.A. Hartl, and G. Bosco. 2012. Condensin II promotes the formation of chromosome territories by inducing axial compaction of polyploid interphase chromosomes. PLoS Genet. 8:e1002873. doi: [10.1371/journal.pgen.1002873].

Bintu, B., L.J. Mateo, J.H. Su, N.A. Sinnott-Armstrong, M. Parker, S. Kinrot, K. Yamaya, A.N. Boettiger, and X. Zhuang. 2018. Super-resolution chromatin tracing reveals domains and cooperative interactions in single cells. Science. 362:eaau1783. doi: [10.1126/science.aau1783].

Bonnet-Garnier, A., P. Feuerstein, M. Chebrout, R. Fleurot, H.U. Jan, P. Debey, and N. Beaujean. 2012. Genome organization and epigenetic marks in mouse germinal vesicle oocytes. Int. J. Dev. Biol. 56:877–887. doi: [10.1387/ijdb.120149ab].

Borsos, M., and M.E. Torres-Padilla. 2016. Building up the nucleus: nuclear organization in the establishment of totipotency and pluripotency during mammalian development. Genes Dev. 30:611–621. doi: [10.1101/gad.273805.115].

Brunner, A., N.R. Morero, W. Zhang, M.J. Hossain, M. Lampe, H. Pflaumer, A. Halavatyi, J.M. Peters, K.S. Beckwith, and J. Ellenberg. 2025. Quantitative imaging of loop extruders rebuilding interphase genome architecture after mitosis. J. Cell Biol. 224:e202405169. doi: [10.1083/jcb.202405169].

Chen, H., D.D. Hughes, T.A. Chan, J.W. Sedat, and D.A. Agard. 1996. IVE (Image Visualization Environment): a software platform for all three-dimensional microscopy applications. J Struct Biol. 116:56–60. doi: [10.1006/jsbi.1996.0010].

Cremer, M., K. Brandstetter, A. Maiser, S.S.P. Rao, V.J. Schmid, M. Guirao-Ortiz, N. Mitra, S. Mamberti, K.N. Klein, D.M. Gilbert, H. Leonhardt, M.C. Cardoso, E.L. Aiden, H. Harz, and T. Cremer. 2020. Cohesin depleted cells rebuild functional nuclear compartments after endomitosis. Nat Commun. 11:6146. doi: [10.1038/s41467-020-19876-6].

Cremer, M., K. Küpper, B. Wagler, L. Wizelman, J. von Hase, Y. Weiland, L. Kreja, J. Diebold, M.R. Speicher, and T. Cremer. 2003. Inheritance of gene density-related higher order chromatin arrangements in normal and tumor cell nuclei. J. Cell Biol. 162:809–820. doi: [10.1083/jcb.200304096].

Cremer, T., and M. Cremer. 2010. Chromosome territories. Cold Spring Harb Perspect Biol. 2:a003889. doi: [10.1101/cshperspect.a003889].

Finn, E.H., G. Pegoraro, H.B. Brandão, A.L. Valton, M.E. Oomen, J. Dekker, L. Mirny, and T. Misteli. 2019. Extensive Heterogeneity and Intrinsic Variation in Spatial Genome Organization. Cell. 176:1502–1515.e1510. doi: [10.1016/j.cell.2019.01.020].

Fisher, N.I. 1995. Statistical analysis of circular data. Cambridge University Press.

Gabriele, M., H.B. Brandão, S. Grosse-Holz, A. Jha, G.M. Dailey, C. Cattoglio, T.S. Hsieh, L. Mirny, C. Zechner, and A.S. Hansen. 2022. Dynamics of CTCF-and cohesin-mediated chromatin looping revealed by live-cell imaging. Science. 376:496–501. doi: [10.1126/science.abn6583].

Gandhi, R., P.J. Gillespie, and T. Hirano. 2006. Human Wapl is a cohesin-binding protein that promotes sister-chromatid resolution in mitotic prophase. Curr. Biol. 16:2406–2417. doi: [10.1016/j.cub.2006.10.061].

Gerlich, D., T. Hirota, B. Koch, J.M. Peters, and J. Ellenberg. 2006. Condensin I stabilizes chromosomes mechanically through a dynamic interaction in live cells. Curr. Biol. 16:333–344. doi: [10.1016/j.cub.2005.12.040].

Gibcus, J.H., K. Samejima, A. Goloborodko, I. Samejima, N. Naumova, J. Nuebler, M.T. Kanemaki, L. Xie, J.R. Paulson, W.C. Earnshaw, L.A. Mirny, and J. Dekker. 2018. A pathway for mitotic chromosome formation. Science. 359:eaao6135. doi: [10.1126/science.aao6135].

Green, L.C., P. Kalitsis, T.M. Chang, M. Cipetic, J.H. Kim, O. Marshall, L. Turnbull, C.B. Whitchurch, P. Vagnarelli, K. Samejima, W.C. Earnshaw, K.H. Choo, and D.F. Hudson. 2012. Contrasting roles of condensin I and condensin II in mitotic chromosome formation. J. Cell Sci. 125:1591–1604. doi: [10.1242/jcs.097790].

Haarhuis, J.H., A.M. Elbatsh, B. van den Broek, D. Camps, H. Erkan, K. Jalink, R.H. Medema, and B.D. Rowland. 2013. WAPL-mediated removal of cohesin protects against segregation errors and aneuploidy. Curr. Biol. 23:2071–2077. doi: [10.1016/j.cub.2013.09.003].

Hara, Y., M. Iwabuchi, K. Ohsumi, and A. Kimura. 2013. Intranuclear DNA density affects chromosome condensation in metazoans. Mol Biol Cell. 24:2442–2453. doi: [10.1091/mbc.E13-01-0043].

Hartl, T.A., H.F. Smith, and G. Bosco. 2008. Chromosome alignment and transvection are antagonized by condensin II. Science. 322:1384–1387. doi: [10.1126/science.1164216].

Hauf, S., E. Roitinger, B. Koch, C.M. Dittrich, K. Mechtler, and J.M. Peters. 2005. Dissociation of cohesin from chromosome arms and loss of arm cohesion during early mitosis depends on phosphorylation of SA2. PLoS Biol. 3:e69. doi: [10.1371/journal.pbio.0030069].

Hirano, T. 2025. Mitotic genome folding. J. Cell Biol. 224:e202504075. doi: [10.1083/jcb.202504075].

Hirota, T., D. Gerlich, B. Koch, J. Ellenberg, and J.M. Peters. 2004. Distinct functions of condensin I and II in mitotic chromosome assembly. J. Cell Sci. 117:6435–6445. doi: [10.1242/jcs.01604].

Hoencamp, C., O. Dudchenko, A.M.O. Elbatsh, S. Brahmachari, J.A. Raaijmakers, T. van Schaik, Á. Sedeño Cacciatore, V.G. Contessoto, R. van Heesbeen, B. van den Broek, A.N. Mhaskar, H. Teunissen, B.G. St Hilaire, D. Weisz, A.D. Omer, M. Pham, Z. Colaric, Z. Yang, S.S.P. Rao, N. Mitra, C. Lui, W. Yao, R. Khan, L.L. Moroz, A. Kohn, J. St Leger, A. Mena, K. Holcroft, M.C. Gambetta, F. Lim, E. Farley, N. Stein, A. Haddad, D. Chauss, A.S. Mutlu, M.C. Wang, N.D. Young, E. Hildebrandt, H.H. Cheng, C.J. Knight, T.L.U. Burnham, K.A. Hovel, A.J. Beel, P.J. Mattei, R.D. Kornberg, W.C. Warren, G. Cary, J.L. Gómez-Skarmeta, V. Hinman, K. Lindblad-Toh, F. Di Palma, K. Maeshima, A.S. Multani, S. Pathak, L. Nel-Themaat, R.R. Behringer, P. Kaur, R.H. Medema, B. van Steensel, E. de Wit, J.N. Onuchic, M. Di Pierro, E. Lieberman Aiden, and B.D. Rowland. 2021. 3D genomics across the tree of life reveals condensin II as a determinant of architecture type. Science. 372:984–989. doi: [10.1126/science.abe2218].

Iida, S., S. Ide, S. Tamura, M. Sasai, T. Tani, T. Goto, M. Shribak, and K. Maeshima. 2024. Orientation-independent-DIC imaging reveals that a transient rise in depletion attraction contributes to mitotic chromosome condensation. Proc Natl Acad Sci U S A. 121:e2403153121. doi: [10.1073/pnas.2403153121].

Isenhart, R., S.C. Nguyen, L. Rosin, W. Cao, P. Walsh, H. Muzaffar, C.E. Ellison, and E.F. Joyce. 2025. The length and strength of compartmental interactions are modulated by condensin II activity. PLoS Genet. 21:e1011724. doi: [10.1371/journal.pgen.1011724].

Kinoshita, K., Y. Tsubota, S. Tane, Y. Aizawa, R. Sakata, K. Takeuchi, K. Shintomi, T. Nishiyama, and T. Hirano. 2022. A loop extrusion-independent mechanism contributes to condensin I-mediated chromosome shaping. J. Cell Biol. 221:e202109016. doi: [10.1083/jcb.202109016].

Kondo, T., and A. Kimura. 2019. Choice between 1-and 2-furrow cytokinesis in Caenorhabditis elegans embryos with tripolar spindles. Mol Biol Cell. 30:2065–2075. doi: [10.1091/mbc.E19-01-0075].

Kyogoku, H., and T.S. Kitajima. 2023. The large cytoplasmic volume of oocyte. J Reprod Dev. 69:1–9. doi: [10.1262/jrd.2022-101].

Lee, J., S. Ogushi, M. Saitou, and T. Hirano. 2011. Condensins I and II are essential for construction of bivalent chromosomes in mouse oocytes. Mol Biol Cell. 22:3465–3477. doi: [10.1091/mbc.E11-05-0423].

Losada, A., M. Hirano, and T. Hirano. 2002. Cohesin release is required for sister chromatid resolution, but not for condensin-mediated compaction, at the onset of mitosis. Genes Dev. 16:3004–3016. doi: [10.1101/gad.249202].

Luppino, J.M., D.S. Park, S.C. Nguyen, Y. Lan, Z. Xu, R. Yunker, and E.F. Joyce. 2020. Cohesin promotes stochastic domain intermingling to ensure proper regulation of boundary-proximal genes. Nat. Genet. 52:840–848. doi: [10.1038/s41588-020-0647-9].

Mardia, K.V. 1967. A Non-Parametric Test for the Bivariate Two-Sample Location Problem. J. R. Stat. Soc. Ser. B (Stat. Methodol.). 29:320–342. doi: [10.1111/j.2517-6161.1967.tb00699.x].

Matthews, N.E., and R. White. 2019. Chromatin Architecture in the Fly: Living without CTCF/Cohesin Loop Extrusion?: Alternating Chromatin States Provide a Basis for Domain Architecture in Drosophila. Bioessays. 41:e1900048. doi: [10.1002/bies.201900048].

Nakaya, M., H. Tanabe, S. Takamatsu, M. Hosokawa, and T. Mitani. 2017. Visualization of the spatial arrangement of nuclear organization using three-dimensional fluorescence in situ hybridization in early mouse embryos: A new “EASI-FISH chamber glass” for mammalian embryos. J Reprod Dev. 63:167–174. doi: [10.1262/jrd.2016-172].

Natsume, T., T. Kiyomitsu, Y. Saga, and M.T. Kanemaki. 2016. Rapid Protein Depletion in Human Cells by Auxin-Inducible Degron Tagging with Short Homology Donors. Cell Rep. 15:210–218. doi: [10.1016/j.celrep.2016.03.001].

Nishide, K., and T. Hirano. 2014. Overlapping and non-overlapping functions of condensins I and II in neural stem cell divisions. PLoS Genet. 10:e1004847. doi: [10.1371/journal.pgen.1004847].

Ono, T., Y. Fang, D.L. Spector, and T. Hirano. 2004. Spatial and temporal regulation of Condensins I and II in mitotic chromosome assembly in human cells. Mol Biol Cell. 15:3296–3308. doi: [10.1091/mbc.E04-03-0242].

Ono, T., A. Losada, M. Hirano, M.P. Myers, A.F. Neuwald, and T. Hirano. 2003. Differential contributions of condensin I and condensin II to mitotic chromosome architecture in vertebrate cells. Cell. 115:109–121. doi:

Ono, T., C. Sakamoto, M. Nakao, N. Saitoh, and T. Hirano. 2017. Condensin II plays an essential role in reversible assembly of mitotic chromosomes in situ. Mol Biol Cell. 28:2875–2886. doi: [10.1091/mbc.E17-04-0252].

Ono, T., D. Yamashita, and T. Hirano. 2013. Condensin II initiates sister chromatid resolution during S phase. J. Cell Biol. 200:429–441. doi: [10.1083/jcb.201208008].

Paulson, J.R., D.F. Hudson, F. Cisneros-Soberanis, and W.C. Earnshaw. 2021. Mitotic chromosomes. Semin. Cell Dev. Biol. 117:7–29. doi: [10.1016/j.semcdb.2021.03.014].

Perea-Resa, C., L. Bury, I.M. Cheeseman, and M.D. Blower. 2020. Cohesin Removal Reprograms Gene Expression upon Mitotic Entry. Mol. Cell. 78:127–140.e127. doi: [10.1016/j.molcel.2020.01.023].

Perpelescu, M., N. Nozaki, C. Obuse, H. Yang, and K. Yoda. 2009. Active establishment of centromeric CENP-A chromatin by RSF complex. J. Cell Biol. 185:397–407. doi: [10.1083/jcb.200903088].

Rao, S.S.P., S.C. Huang, B. Glenn St Hilaire, J.M. Engreitz, E.M. Perez, K.R. Kieffer-Kwon, A.L. Sanborn, S.E. Johnstone, G.D. Bascom, I.D. Bochkov, X. Huang, M.S. Shamim, J. Shin, D. Turner, Z. Ye, A.D. Omer, J.T. Robinson, T. Schlick, B.E. Bernstein, R. Casellas, E.S. Lander, and E.L. Aiden. 2017. Cohesin Loss Eliminates All Loop Domains. Cell. 171:305–320.e324. doi: [10.1016/j.cell.2017.09.026].

Rosin, L.F., S.C. Nguyen, and E.F. Joyce. 2018. Condensin II drives large-scale folding and spatial partitioning of interphase chromosomes in Drosophila nuclei. PLoS Genet. 14:e1007393. doi: [10.1371/journal.pgen.1007393].

Rowley, M.J., M.H. Nichols, X. Lyu, M. Ando-Kuri, I.S.M. Rivera, K. Hermetz, P. Wang, Y. Ruan, and V.G. Corces. 2017. Evolutionarily Conserved Principles Predict 3D Chromatin Organization. Mol. Cell. 67:837–852.e837. doi: [10.1016/j.molcel.2017.07.022].

Sakamoto, T., Y. Sakamoto, S. Grob, D. Slane, T. Yamashita, N. Ito, Y. Oko, T. Sugiyama, T. Higaki, S. Hasezawa, M. Tanaka, A. Matsui, M. Seki, T. Suzuki, U. Grossniklaus, and S. Matsunaga. 2022. Two-step regulation of centromere distribution by condensin II and the nuclear envelope proteins. Nat Plants. 8:940–953. doi: [10.1038/s41477-022-01200-3].

Samejima, K., J.H. Gibcus, S. Abraham, F. Cisneros-Soberanis, I. Samejima, A.J. Beckett, N. Pučeková, M.A. Abad, C. Spanos, B. Medina-Pritchard, J.R. Paulson, L. Xie, A.A. Jeyaprakash, I.A. Prior, L.A. Mirny, J. Dekker, A. Goloborodko, and W.C. Earnshaw. 2025. Rules of engagement for condensins and cohesins guide mitotic chromosome formation. Science. 388:eadq1709. doi: [10.1126/science.adq1709].

Schwarzer, W., N. Abdennur, A. Goloborodko, A. Pekowska, G. Fudenberg, Y. Loe-Mie, N.A. Fonseca, W. Huber, C.H. Haering, L. Mirny, and F. Spitz. 2017. Two independent modes of chromatin organization revealed by cohesin removal. Nature. 551:51–56. doi: [10.1038/nature24281].

Shintomi, K., and T. Hirano. 2011. The relative ratio of condensin I to II determines chromosome shapes. Genes Dev. 25:1464–1469. doi: [10.1101/gad.2060311].

Sun, Y., X. Xu, W. Zhao, Y. Zhang, K. Chen, Y. Li, X. Wang, M. Zhang, B. Xue, W. Yu, Y. Hou, C. Wang, W. Xie, C. Li, D. Kong, S. Wang, and Y. Sun. 2023. RAD21 is the core subunit of the cohesin complex involved in directing genome organization. Genome Biol. 24:155. doi: [10.1186/s13059-023-02982-1].

Takagi, M., T. Ono, T. Natsume, C. Sakamoto, M. Nakao, N. Saitoh, M.T. Kanemaki, T. Hirano, and N. Imamoto. 2018. Ki-67 and condensins support the integrity of mitotic chromosomes through distinct mechanisms. J. Cell Sci. 131:jcs212092. doi: [10.1242/jcs.212092].

Tanabe, H., F.A. Habermann, I. Solovei, M. Cremer, and T. Cremer. 2002. Non-random radial arrangements of interphase chromosome territories: evolutionary considerations and functional implications. Mutat. Res. 504:37–45. doi: [10.1016/s0027-5107(02)00077-5].

Vassilev, L.T., C. Tovar, S. Chen, D. Knezevic, X. Zhao, H. Sun, D.C. Heimbrook, and L. Chen. 2006. Selective small-molecule inhibitor reveals critical mitotic functions of human CDK1. Proc Natl Acad Sci U S A. 103:10660–10665. doi: [10.1073/pnas.0600447103].

Wesley, C.C., S. Mishra, and D.L. Levy. 2020. Organelle size scaling over embryonic development. Wiley Interdiscip Rev Dev Biol. 9:e376. doi: [10.1002/wdev.376].

Wutz, G., C. Várnai, K. Nagasaka, D.A. Cisneros, R.R. Stocsits, W. Tang, S. Schoenfelder, G. Jessberger, M. Muhar, M.J. Hossain, N. Walther, B. Koch, M. Kueblbeck, J. Ellenberg, J. Zuber, P. Fraser, and J.M. Peters. 2017. Topologically associating domains and chromatin loops depend on cohesin and are regulated by CTCF, WAPL, and PDS5 proteins. EMBO J. 36:3573–3599. doi: [10.15252/embj.201798004].

